# A female-biased gene expression signature of dominance in cooperatively breeding meerkats

**DOI:** 10.1101/2023.12.01.569325

**Authors:** C. Ryan Campbell, Marta Manser, Mari Shiratori, Kelly Williams, Luis Barreiro, Tim Clutton-Brock, Jenny Tung

## Abstract

Dominance is a primary determinant of social dynamics and resource access in social animals. Recent studies show that dominance is also reflected in the gene regulatory profiles of peripheral immune cells. However, the strength and direction of this relationship differs across the species and sex combinations investigated, potentially due to variation in the predictors and energetic consequences of dominant status. To test this possibility, we investigated the association between social status and gene expression in the blood of wild meerkats (*Suricata suricatta*; n=113 unique individuals), including in response to lipopolysaccharide, Gardiquimod, and glucocorticoid stimulation. Meerkats are cooperatively breeding social carnivores in which breeding females physically outcompete other females to suppress reproduction, resulting in high reproductive skew. They therefore present an opportunity to disentangle the effects of social dominance from those of sex *per se*. We identify a sex-specific signature of dominance, including 1,045 differentially expressed genes in females but none in males. Dominant females exhibit elevated activity in innate immune pathways and an exacerbated response to LPS challenge. In this respect, female meerkats resemble wild male baboons (for which similar data are available), where physical competition is also central to determining rank hierarchies and mating effort is high. However, they differ from female primates in which social status is nepotistically determined. Our results support the hypothesis that the gene regulatory signature of social status depends on the determinants and energetic costs of social dominance. They also support potential life history trade-offs between investment in reproduction versus somatic maintenance.

## Introduction

In species that live in stable groups, social dominance often predicts access to food, mates, social partners, and other monopolizable resources (Clutton-Brock, 1988; Clutton-Brock, 2021; Krause & Ruxton, 2002; Rubenstein, 1978; Sterck et al., 1997; Wilson, 2000). However, there is substantial variation in how animals attain and maintain dominance, both between and within species (e.g., between sexes). While some social systems are shaped by familial (nepotistic) inheritance, such that individuals attain a rank position similar to their relatives (e.g., female Japanese macaques: Kawai (1958); Kawamura (1958); female yellow baboons: Hausfater et al. (1982); female spotted hyenas: Holekamp and Smale (1991); female geladas: Le Roux et al. (2011)), others are based on individual characteristics such as physical condition or fighting ability (e.g., male red deer: Clutton-Brock et al. (1982); male yellow baboons: Alberts et al. (2003); male bottlenose dolphins: Samuels and Gifford (1997)). Dominance status also varies in temporal stability and in the behavioral expression of dominance. For example, in the macaque radiation, rhesus macaque females enforce dominance via regular harassment directed down a predictable linear hierarchy (Bernstein et al., 1974). In contrast, in closely related crested macaques, agonistic interactions are bidirectional, rarely involve physical threat or contact, and are often followed by conciliatory behavior (Duboscq et al., 2013). Consequently, while high status is generally fitness-enhancing, its physiological costs and benefits are expected to vary.

Variation in the costs and benefits of high status is perhaps best supported by studies of the relationship between dominance and glucocorticoid levels, a key hormonal marker of both energetic and psychosocial stress (Abbott et al., 2003; Beehner & Bergman, 2017; Cavigelli & Caruso, 2015; Creel, 2005). Although experimental studies in lab models generally link low status to high glucocorticoid levels and/or glucocorticoid resistance (e.g., Kohn et al. (2016); Razzoli et al. (2018); Willard and Shively (2012); reviewed in: Beehner and Bergman (2017); Cavigelli and Caruso (2015)), research across a wider range of species, including in natural populations, paints a more complex picture. Higher glucocorticoids correlate with low status in some cases, especially when dominance is aggressively enforced or opportunities for social support are lacking (Abbott et al., 2003). However, high glucocorticoid levels are linked to *high* status in other settings (in mammals, most often in males), potentially because of the high energetic demands of achieving high rank, defending against challengers, or investing in reproductive opportunities (Creel et al., 1997; Fichtel et al., 2007; Gesquiere et al., 2011; Muller et al., 2021; Muller & Wrangham, 2004; Schoof & Jack, 2013). In male chimpanzees, for example, the positive correlation between high status and glucocorticoid production appears to be mediated by rates of aggressive behavior directed by dominant individuals towards social subordinates (Muller et al., 2021; Muller & Wrangham, 2004). In cases where high status is associated with high glucocorticoid levels, high status has also been linked to other, presumably costly, outcomes. For instance, in male baboons in the Amboseli study population, high status predicts high glucocorticoid levels, accelerated epigenetic age, and a moderately elevated mortality risk (Anderson et al., 2021; Campos et al., 2021; Campos et al., 2020).

Recent studies have begun linking variation in social status to gene regulation, producing a novel molecular source of insight into the potential costs and benefits of dominance. While the earliest such studies focused on gene expression changes involved in the initial transition to dominance, especially in the brain (e.g., the emergence of honeybee queens Evans & Wheeler, 2000; Grozinger et al., 2003; Maruska et al., 2013), more recent work has sought to understand how the different experiences of high versus low status individuals translate to changes in gene regulation in the periphery (Cole, 2014; Simons & Tung, 2019). Strong support for status-related effects comes from lab studies of mice, in which low status animals and those subjected to repeated social defeat show substantially altered peripheral blood mononuclear cell (PBMC) gene expression (Lee et al., 2022b; Pizarro et al., 2004; Powell et al., 2013). In primates, experimental manipulations of group hierarchies in rhesus macaques alter not only baseline gene expression, but also the response to immune stimulation (Sanz et al., 2020; Snyder-Mackler et al., 2016; Snyder-Mackler et al., 2019). In general—and in agreement with correlational studies in humans (Cole, 2019; Miller et al., 2009; Murray et al., 2019; Powell et al., 2013)—these studies suggest that low social status is typically associated with elevated activity of pro-inflammatory pathways. Conversely, high status is frequently linked to increased activity of genes involved in interferon-induced antiviral defense.

However, the settings in which links between social status and gene regulation are most pronounced have not been systematically explored. Recent studies in wild baboons indicate that there may be substantial variation in the strength and direction of social status associations with gene expression in immune cells (Anderson et al., 2022; Lea et al., 2018). For example, while low-ranking baboon females show a pattern of pro-inflammatory/anti-viral polarization similar to that reported for socially stressed humans and captive rhesus macaques (Murray et al., 2019; Snyder-Mackler et al., 2016), this pattern is reversed for baboon males from the same study population. Viewed through the lens of gene expression, high status males therefore resemble low status females, and vice-versa (Anderson et al., 2022; Lea et al., 2018). One possible explanation for this contrast involves the distinct dynamics of status competition in male versus female baboons. While male baboons physically compete to establish dominance, young females typically insert in the status hierarchy directly below their mothers (Hausfater et al., 1982; Lea et al., 2014). Additionally, because high-ranking male baboons are able to monopolize mating opportunities whereas all adult females are typically able to breed, reproductive skew is greater in males than in females (Alberts et al., 2006; Lukas & Clutton-Brock, 2014), consistent with typical observations in polygynous or polygynandrous mammals (Clutton-Brock, 1988). Differences in the level of physical competition for social dominance and status-related variance in reproductive investment may therefore play a central role in understanding the gene regulatory signature of social status.

Testing this hypothesis requires separating patterns of competition and reproductive investment from the effects of sex *per se.* Although many mammal societies follow the pattern in baboons (i.e., more intense competition and energetic investment in mating opportunities in males), the typical pattern of sex differences in the correlates of dominance status is reversed in some mammalian species (e.g., Clutton-Brock and Huchard (2013); Lukas and Clutton-Brock (2014); see also Emlen and Wrege (2004); Goymann et al. (2004); Oring and Lank (1986) for work outside of mammals). In cooperatively breeding meerkats (*Suricata suricatta*), for instance, reproduction is concentrated in one dominant breeding female per group, who defends her position via regular aggression directed at competitors, including targeted eviction and infanticide if subordinate animals attempt to breed (Clutton-Brock et al., 2001; Kutsukake & Clutton-Brock, 2006; Young et al., 2006). Because subordinates of both sexes assist in rearing young, dominant females can also breed multiple times a year. As a result, reproductive skew in female meerkats can be extreme: in one example, a successful dominant female reared 72 offspring during her lifetime, whereas most subordinates produce no surviving offspring (Clutton-Brock et al., 2006; Hodge et al., 2008). Dominance in males also leads to skewed reproduction, as dominant males also father most pups born to the dominant female in their groups (Griffin et al., 2003; Hodge et al., 2008; Spong et al., 2008). However, because the tenure of dominant males is typically shorter than that of dominant females, and because they tend to have shorter breeding lifespans, variance in lifetime breeding success is lower than for females and competition for breeding opportunities is more frequent and more intense in females than in males (Clutton-Brock et al., 2001; Clutton-Brock et al., 2006; Clutton-Brock & Manser, 2016; Duncan et al., 2023; Hodge et al., 2008; Lukas & Clutton-Brock, 2014; Spong et al., 2008). Attainment of dominant status is also body mass (i.e., condition)-dependent in females, but not in males (Duncan et al., 2018; Spong et al., 2008). Dominance is associated with higher cortisol levels in both sexes (Carlson et al., 2004), but female meerkats exhibit higher parasite load than males (Smyth & Drea, 2016), and dominant females (but not males) have higher androgen levels than their same-sex subordinates (Carlson et al., 2004; Drea & Davies, 2022; Drea et al., 2021).

The unusual pattern of sex differences in meerkats therefore provides an ideal test case for investigating the patterns responsible for associations between social status and immune gene expression. To do so here, we investigate the association between social status and gene regulation in wild meerkats of both sexes. We focus on gene expression patterns in circulating peripheral immune cells, which are accessible using minimally invasive methods, facilitate comparisons to previous work in nonhuman primates, and capture aspects of immune function that may trade-off against dominance-related investment in body condition or reproductive effort. First, we ask if dominance status predicts immune gene expression, and whether these signatures differ by sex. Second, we investigate whether social status is also associated with the immune gene expression response to pathogen stimuli, as it is in baboons and rhesus macaques (Lea et al., 2018; Snyder-Mackler et al., 2016), and consistent with status-related differences in parasite load previously described for this species (Dantzer et al., 2017a; Smyth & Drea, 2016). Third, we assess whether status-related gene expression differences in females are explained by age, body mass, or reproductive state. Finally, we place our results for meerkats in context by comparing the pathways associated with status in this population to those reported for two, non-cooperatively breeding primates in which physical competition for rank and reproductive skew is male-biased (Alberts et al., 2006; Anderson et al., 2022; Lea et al., 2018; Lukas & Clutton-Brock, 2014; Snyder-Mackler et al., 2016; Widdig et al., 2004; Widdig et al., 2016). Together, our work shows how diverse social structures shape the gene regulatory signature of dominance and contribute to an emerging understanding of the processes that link social status to its molecular correlates.

## Methods

### Field site and sampling

This study was conducted on 129 wild meerkats (69 males and 60 females), members of 15 study groups monitored by the Kalahari Meerkat Project at the Kuruman River Reserve, Northern Cape Province, South Africa, between August 2017 and September 2020 (Table S1). These groups were visited 3 – 7 days per week to collect demographic, life history, and behavioral data, including agonistic interactions used to infer dominance status (Clutton-Brock et al., 1998). The majority of blood samples used in this study were collected cross-sectionally during scheduled biannual or annual draws (n=237 blood samples from 129 unique individuals). However, we also targeted newly dominant individuals for longitudinal sampling, with blood collected (i) as close to the date of their transition from subordinate to dominant status as possible, (ii) four weeks after their dominance transition, and (iii) four months after their initial transition (assuming the animal retained dominant breeding status). We simultaneously sampled a subordinate member of the same group and a subordinate member of a different group for comparison, allowing us to control for temporal, seasonal, and group effects. 98 total blood draws were collected based on this longitudinal design. The meerkats in the study population were habituated to humans and are captured by hand. Meerkats were anaesthetized using 4% isoflurane (Isofor, Safe Line Pharmaceuticals, Johannesburg, South Africa) mixed with oxygen administered via a gas mask attached to a portable vaporizer. After full sedation, isoflurane dosage was lowered to 1–2% and a blood sample was obtained from the jugular vein using a 25 G needle and 2-ml syringe (as in Carlson et al., 2004; Dantzer et al., 2017a; Drea et al., 2021).

To assess dominance status as well as pregnancy we drew on regular behavioral and demographic observations, as described in Clutton-Brock et al. (1998). Dominant animals routinely displace subordinates, scent-mark more often, and are typically older and, in the case of females, heavier than their groupmates (Figure S1). They are also more likely to be reproductively active, although not exclusively so, as subordinates sometimes become pregnant, although rarely successfully raise young. Pregnancy was assessed via observations of physical expansion of the abdomen, which becomes visually obvious roughly halfway through the 70-day gestation period (Clutton-Brock et al., 1998). Fifteen samples were collected when females were likely pregnant (13% of the 119 samples). Because we avoided sampling visibly pregnant female meerkats, 93% of these samples were collected in the first half of pregnancy (11 – 39 days gestation). Nine of these 15 cases were samples collected from dominant females and 6 were from subordinate individuals, but consistent with high reproductive skew in female meerkats, only one pregnancy from a subordinate female resulted in live offspring that survived more than one week.

Finally, to assess body mass, study subjects were weighed multiple times daily using electronic balances (Clutton-Brock et al., 1998). For each capture effort, body mass was determined by taking the focal study subject’s average measure of body mass across all measures collected within 30 days of the capture date, excluding 2.3% of measurements identified as outliers (Grubbs outlier test of 10 consecutive weights, p<0.05). Age at sampling (mean = 2.1 years ± 1.44 years; range = 0.96-10.4 years; Table S1) was based on long-term observations and was known to within a few days’ error for 97% of individuals in the sample.

### Cell challenge and RNA-seq data generation

We purified PBMCs from ∼1 mL of blood per blood draw via density gradient centrifugation, typically within 3 – 4 hours of the original blood draw. For each blood sample, we plated 200,000 PBMCs in a 200 µL cell suspension into each of four tissue culture wells containing 20 µL cell culture media (RPMI; 10% FBS; 1% penicillin-streptomycin), for a total volume of 220 µL. Purified PBMCs from each sample were cultured for 4 hours in (i) media only (control condition), (ii) media with 10 ng/mL lipopolysaccharide (LPS from the *E. coli* O111:B4 strain), to mimic bacterial exposure; (iii) media plus 1.0 μg/mL Gardiquimod (Gard), which activates Toll-like receptor 7 signaling; or (iv) media plus 1.0 μM Dexamethasone (Dex), a synthetic glucocorticoid. The cells were then incubated in parallel for 4 hours (37° C and 5% CO_2_), washed with 1× PBS, lysed, and stored immediately at -80° C until library preparation.

We prepared RNA-sequencing libraries for each sample (control, Dex-challenged, Gard-challenged, LPS-challenged) by purifying mRNA using the miRNeasy Mini Kit (Qiagen). Libraries were generated following the Transposase Mediated 3′ RNAseq (TM3’seq) protocol (Pallares et al., 2020) with an input of 50 ng total RNA. This protocol consists of first-strand cDNA synthesis, amplification, tagmentation, and a final library amplification step. It generates 3’ tags to represent transcripts as opposed to conventional full transcript RNA-seq, reducing the read depth required to represent the transcriptome (Pallares et al., 2020). All libraries were sequenced on an Illumina NovaSeq 6000 at the University of Chicago’s Genomics Facility using 100 bp single-end reads.

### Data analysis

TM-3’-seq data were first filtered with trimmomatic (version 0.38, Bolger et al., 2014) to remove adapter sequence and stretches of low-quality bases, as well as reads <20 bp following trimming. Remaining trimmed reads were mapped to the meerkat genome (GCF_006229205.1, https://www.ncbi.nlm.nih.gov/datasets/genome/GCF_006229205.1/) using the STAR two-pass method (version 2.7.10ac, Dobin et al., 2013). Mapped reads were aggregated to the gene level using HTseq (version 2.0, Putri et al., 2022) based on overlap with annotated gene exons. As expected based on the library preparation method we used, mapped reads were highly biased towards the 3’ end of genes (Figure S2). We excluded 199 libraries from further analysis because the number of genes with non-zero read count was low (<10,000, compared to mean=11,690); typically, these libraries also were shallowly sequenced overall. We also removed 11 samples where the distribution of gene expression levels (logCPM) was systematically lower for most genes than observed on average.

To investigate the gene expression signature of treatment (LPS, Gard, and Dex compared to control samples) and dominance status, we focused on genes that were moderately to highly expressed in meerkat PBMCs, based on mean expression across samples (n=6,012 – 6,932 genes with mean log(CPM) > 5, depending on sex-culture condition combination; Table S3). To control for differences in sequencing depth and technical effects of library quality and batch, we normalized the gene count matrix using the function *voom* from the R package limma (version 3.54.1, 2015; Ritchie et al., 2015) and regressed out library batch, number of unique genes, percent of reads in feature, and percent of reads mapped to the genome. We first explored the effect of LPS, Gard, and Dex stimulation in both sexes and in models for males and females separately (see “Treatment-control” models in the Supplementary Methods). Here, we controlled for mean-centered body mass, mean-centered age at sampling, dominance status (represented as a binary variable for dominant versus subordinate), and pregnancy status (represented as a binary variable, in females only) as fixed effect covariates.

These initial models pointed to shared effects of stimulation in both sexes but indicated that dominance status effects were only detectable in females. To test this possibility further, we fit sex-specific models considering data from all treatment conditions together (including condition as a fixed effect in addition to body mass, age, pregnancy status, and dominance status; see “All conditions, sex-specific” model in the Supplementary Methods). Because the results of these analyses again indicated that dominance status effects are detectable only in females, we ran subsequent models for female meerkats only. We first investigated dominance effects in each of four conditions separately (control, LPS-stimulated, Gard-stimulated, and Dex-stimulated; “Condition-specific, female-only” model in the Supplementary Methods). Finally for the subset of genes with evidence for gene expression-dominance status associations (n=735, 10% FDR in any condition), we tested for a link between dominance status and the response to challenge by modeling data from the control condition and the relevant stimulated condition(s), nesting the binary dominance status variable within condition (“Status x treatment interactions, female-only” model in the Supplementary Methods). We again controlled for body mass, age, pregnancy status, and included a fixed effect of condition itself. We then compared the estimated effect sizes for dominance status within the control condition and the stimulated condition.

All models controlled for relatedness within the sample (R package: EMMREML version 3.1, Akdemir & Okeke, 2015). To control for multiple hypothesis testing, we calculated false discovery rates based on comparison of the observed p-value distributions to empirical null distributions generated via permutation (Storey & Tibshirani, 2003). To create the empirical null, we permuted dominance status (or other predictor variables of interest, such as treatment) across samples, with samples from the same capture treated as a block. We then fit the same model as in the real data.

For females followed longitudinally across a dominance transition (n=5 who transitioned, plus n=5 controls sampled at the same time, who did not transition), we focused on 514 genes identified as dominance-associated in the full data set (after removing samples from multiple capture dates for the longitudinally followed individuals: see Supplementary Methods). For each of the 514 genes, for each of the five females, we calculated the within-individual log-fold change of the difference in gene expression levels between samples collected when the female was dominant versus subordinate. If multiple samples were collected from a given individual when she was subordinate or dominant, we randomly chose a single sample to represent each dominance category. Because pregnancy also may affect gene expression (Figure S3), we also required that both the “subordinate” and “dominant” sample for the same individual be matched for reproductive status: either both samples must have been collected when the female was pregnant (n=1), or both samples must have been collected when the female was not pregnant (n=4).

To test for enrichment of pathways and gene sets within dominance-associated genes, we conducted gene set enrichment analysis (GSEA) using the Molecular Signatures Database Hallmark Gene Sets (Liberzon et al., 2015) against the background set of all genes included in our analyses, with the p-value for enrichment estimated based on comparison to GSEA test statistics derived from permuting observed effect sizes across genes. In addition, based on prior work indicating social status-associated polarization of TLR4 signaling pathways (Irwin & Cole, 2011; Lea et al., 2018; Slavich & Cole, 2013; Snyder-Mackler et al., 2016), we annotated a custom gene set of MyD88- and TRIF-dependent genes based on the results of Ramsey et al. (2008). Specifically, we identified genes that were upregulated after immune stimulation in wild-type mice in a MyD88- or TRIF-dependent manner, as determined based on comparison to MyD88 or TRIF knockout mice (Ramsey et al., 2008).

Finally, to compare the gene expression signature of dominance status across female meerkats and nonhuman primates, we drew on previously published results from male and female baboons (Anderson et al., 2022) and female rhesus macaques (Snyder-Mackler et al., 2016). To make comparisons across these four species-sex combinations, we extracted standardized effect sizes for the effect of dominance rank/social status, as reported in each original analysis. We then tested for pathway/gene set enrichment against the background set of genes analyzed for the same sex-species combination. To aid with interpretability, we reverse-coded effect sizes from the baboon data set, where in the original data, high numerical values for rank correspond to low status. Consequently, for all data sets, positive effect sizes correspond to genes in which expression levels are higher in dominant/higher status individuals, and vice-versa.

Statistical analyses were conducted in R (v4.1.2, R Development Core Team, 2021). All analysis code and necessary files are available at https://github.com/cryancampbell/meerkatPaper.

## Results

### A sex-specific signature of breeding status in peripheral blood gene expression patterns

Following quality control, our final data set included RNA-seq profiles from 113 individually recognized meerkats in the Kalahari study population (740 total samples from 52 females and 61 males in 15 social groups, obtained across 200 blood draws; Figure 1; Table S1; mean = 5.87 million ± 5.06 million s.d. reads; Table S2). 59 animals were dominant at the time of capture (31 females and 28 males) and 141 were subordinate at the time of capture. Forty individuals were sampled repeatedly, 15 of whom are represented in our data set as both subordinates and dominants (i.e., were sampled across a dominance transition). Note that we focus on 5 of these individuals for our longitudinal analysis below, based on the set of samples (i) from females; and (ii) for whom both subordinate and dominant samples were collected in the same pregnancy state.

**Figure 1.**
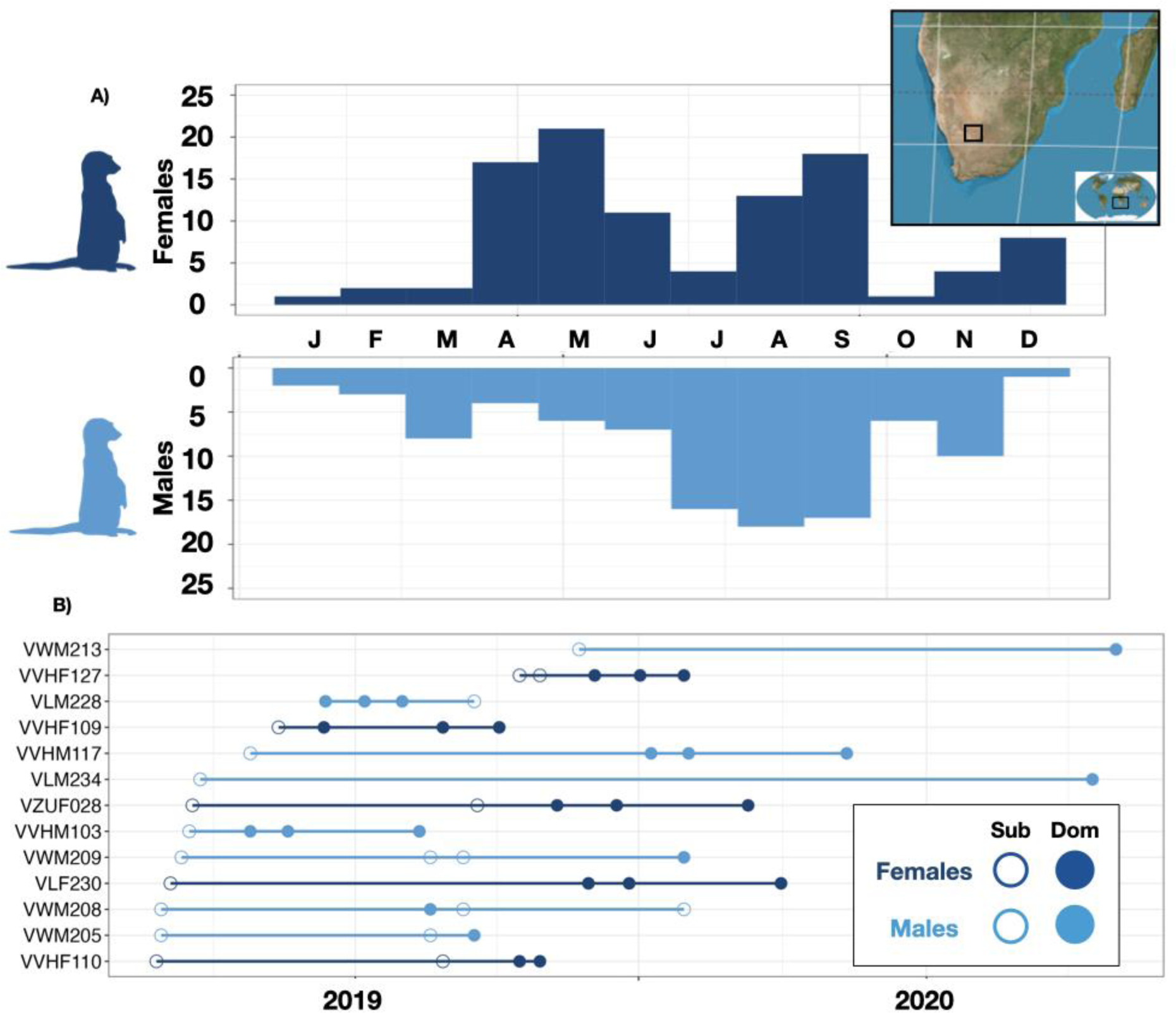
Sampling strategy. A) Sampling by calendar month across the study, broken down by month and sex (females dark blue, males light blue). Inset map shows location of the study population B) Longitudinal sampling for individuals sampled as both dominants and subordinates. Each line represents one individual, and the horizontal axis shows time across multiple years of sample collection. Empty circles represent samples collected when the animal was subordinate and filled circles represent samples collected when the animal was dominant.

We observed a strong gene regulatory response to stimulation in our samples. For all three stimulants (LPS, Gard, and Dex: Figure 2A), control samples separate from stimulated samples along the first and/or second principal component of variation in overall gene expression levels (r_PC1-treatment condition_=-0.91, p=7.80×10^-139^ for LPS, n=6,470 genes; r_PC2-treatment condition_=-0.78, p=3.12×10^-78^ for Gard, n=6,381 genes; r_PC1-treatment condition_=0.66, p=4.46×10^-48^ and r_PC2-treatment condition_=0.64, p=2.04×10^-44^ for Dex, n=6,725 genes; Figure 2B). Responses to all stimulants are highly consistent between males and females (Figure 2C; Figure S4; Table S3), and responsive genes for all three stimulants are strongly enriched in pathways related to innate immunity (e.g., TNFα signaling via NFkB, all p_adj_ < 5 x 10^-4^; inflammatory response, all p_adj_ < 5 x 10^-4^). As expected, stimulation with LPS and Gard, but not Dex, strongly increased the activity of multiple innate immune defense pathways, including IL6 signaling, interferon signaling, and the pro-inflammatory response (Figure 2D).

**Figure 2.**
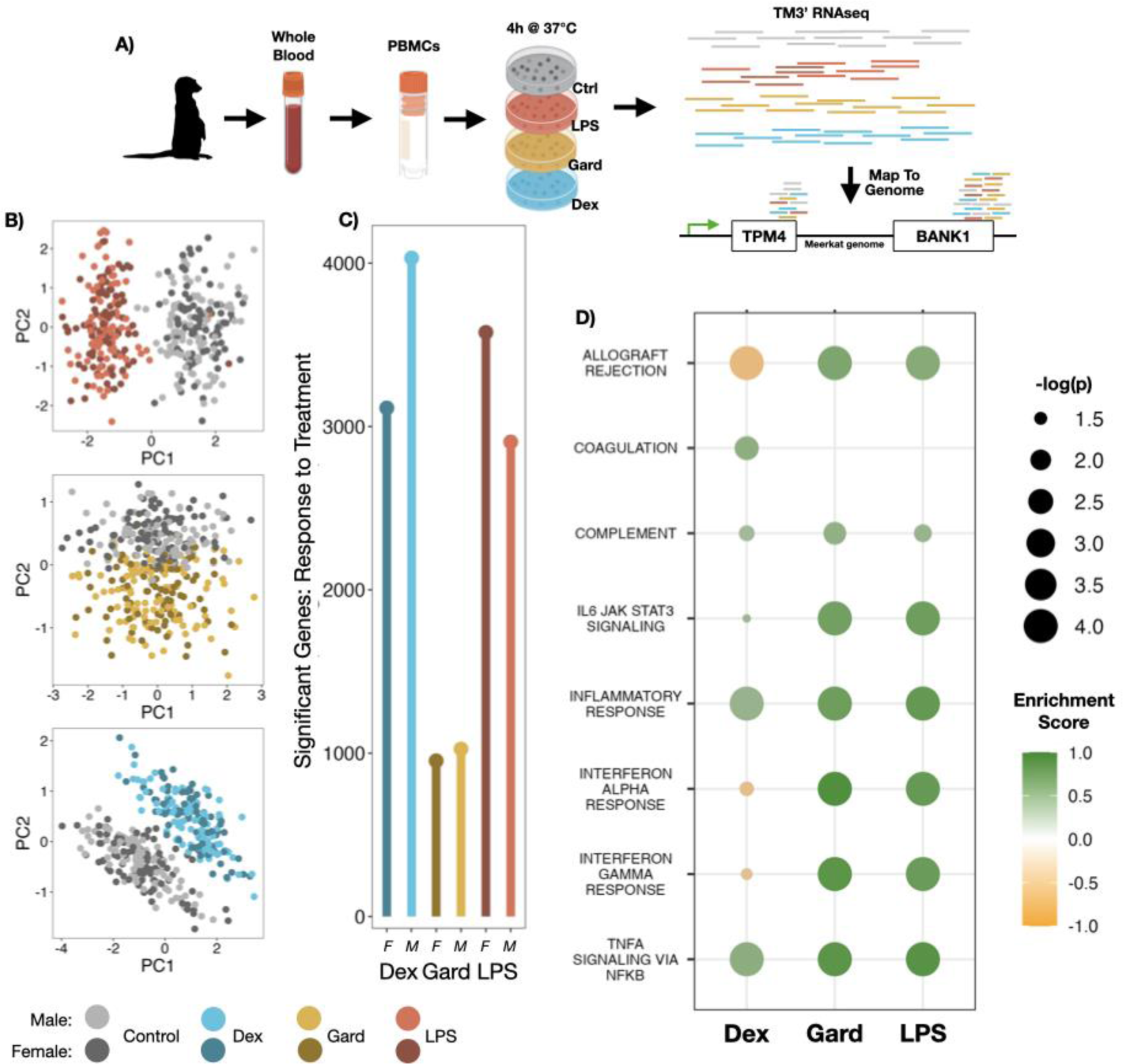
Strong gene expression responses to ex vivo stimulation in meerkat PBMCs. **A)** PBMCs were purified from blood samples obtained during each capture, aliquoted into wells with culture media, and cultured for 4 hours at 37°C under control, LPS-challenged, Gard-challenged, or Dex-challenged conditions. TM-3’seq libraries were prepared from each capture-culture condition combination. **B)** The top principal components of the gene expression data separate each treatment condition (LPS, Gard, and Dex) from the control condition, in both sexes, as supported by **C)** the large number of genes that show a significant response to stimulation (relative to the control condition) for all three conditions, in both males and females. Dark shades = females; light shades = males. **D)** Strong enrichment for genes involved in innate immunity among treatment-responsive genes, especially for LPS and Gard-responsive genes. Circle size is scaled by p-value for enrichment (all plotted points are significant at a nominal p < 0.05) and color represents the enrichment score: green circles are more highly expressed in the treatment condition and orange circles are more highly expressed in the control condition.

In analyses across samples including both sexes (“Treatment-control Models”), nesting dominance status (dominant breeder versus subordinate helper) within sex, our data suggested a strong signature of dominance in female meerkats but not male meerkats. In support of this possibility, a subsequent analysis in females only, across all culture conditions, identified 1,045 dominance status-associated genes (FDR < 10%; 15% of 6,932 analyzed genes, Figure 3A), compared to 0 status-associated genes in males. Dominant females tend to upregulate genes involved in the inflammatory response and NFkB signaling (TNFα signaling via NFkB, enrichment p_adj_ < 5 x 10^-4^ in the LPS challenge condition; inflammatory response, p_adj_ < 5 x 10^-4^ in the LPS challenge condition, Figure 5A) relative to subordinate helpers. Sex differences are unlikely to result from differences in power, as our sample size for males (n=61 unique males; n=359 samples) exceeds our sample size for females (n=52 unique females; n=381 samples). Further, estimates for the dominance effect in males versus females are not well-correlated (Pearson’s R=-0.07, p=1.20×10^-9^, Figure S5). Our results therefore support highly sex-specific effects of dominance on meerkat gene regulation, consistent with previous findings in wild baboon blood cells (analogous to the sample type used here: Anderson et al. (2022); Lea et al. (2018)), cichlid brain (Burmeister et al., 2005), and mouse brain, liver, and spleen (Lee et al., 2022a; Lee et al., 2022c).

**Figure 3.**
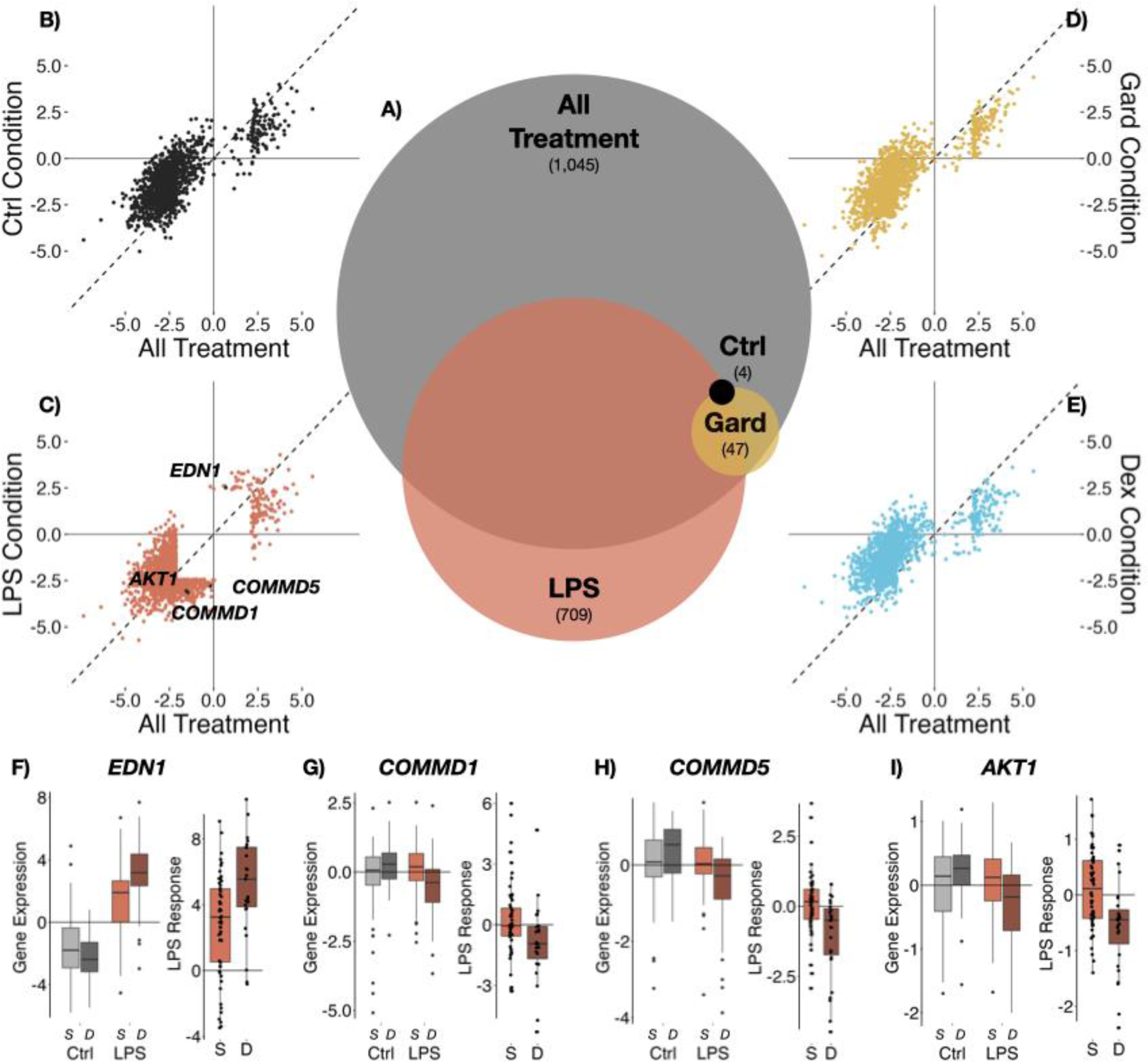
The gene expression signature of dominance status in female meerkats. **A)** Venn diagram of status-associated genes (FDR < 0.10) across each condition-specific model in females, relative to a model including data from all conditions. Dex is not shown here because no genes pass a 10% FDR cut-off in the Dex-specific model. **B-E)** Scatterplot of standardized effect sizes for dominance status in a model including all data (x-axis) versus data from single treatment conditions (y-axis; B: control; C: LPS; D: Gard; E: Dex). Dots correspond to the union set of all genes identified as status-associated in any condition or in the model including all data. **F-H)** Gene expression levels for subordinate (S) and dominant (D) female meerkats in the control condition versus the LPS condition for three immune-related genes (left plot for each panel) in which the response to LPS stimulation systematically differs by social status (right plot for each panel). The difference in response was modeled as an effect of dominance status on within-individual log-foldchange in gene expression after stimulation (EDN1: q=4.0 x 10^-2^; COMMD1: q=4.7 x 10^-2^; COMMD5: q=4.7 x 10^-2^; AKT1: q=4.7 x 10^-2^).

### Treatment condition-dependent effects of female dominance

To identify the conditions in which the dominance signature is most pronounced in females, we modeled gene expression levels as a function of dominance status in control, LPS-challenged, Gard-challenged, and Dex-challenged samples separately (again controlling for body mass, age, and pregnancy status; control: n=96 samples and 6230 genes with log(CPM) > 5; LPS: n=91 samples, 6012 genes; Gard: n=98 samples, 6119 genes; Dex: n=96 samples, 6416 genes; Table S4). Our results indicate that the overall signature of dominance is driven by immune-challenged cells (Figure 3A). While only 4 genes are significantly associated with dominance in the control condition (10% FDR) and no dominance-associated genes are detectable in the Dex-challenged condition, 47 genes and 709 genes are dominance-associated in the Gard and LPS conditions, respectively.

Together, these observations suggest that conditions that stimulate a pro-inflammatory innate immune response magnify differences between dominant and subordinate meerkats (glucocorticoid stimulation may also attenuate them, although the evidence here is less clear: Figure S6). In support of this possibility, dominance status also significantly predicts the *magnitude* of the gene regulatory response to LPS (i.e., the fold-change difference between LPS-challenged samples and control samples for the same individual) for 26 genes (10% FDR, Table S5). These genes include endothelin-1 (*EDN1,* Figure 3F), which is a key component of wound healing and inflammation (Khimji & Rockey, 2010); COMM Domain-containing genes *COMMD1* and *COMMD5* (Figure 3G), which regulate the transcription factor NF-κB (Maine & Burstein, 2007); and the protein kinase *AKT1* (Figure 3H) which regulates the response of macrophages to LPS (Androulidaki et al., 2009).

### Validation of dominance status-related gene expression in longitudinal samples

If females differ in PBMC gene expression because of their dominance status, then females who were sampled first as subordinates and later as dominants should show shifts in gene expression levels that are congruent with our findings above. To test this prediction, we focused on five females who were sampled longitudinally across a transition to dominance. As a control for technical effects of sampling and/or ecological and demographic differences across the elapsed time between these samples, we also compared longitudinal differences in gene expression for five matched females sampled at the same time (± 1 day), but who did *not* undergo a transition (i.e., remained subordinate throughout the same period).

For the 514 gene-condition combinations we identified as status-associated in the cross-sectional analysis (this number reflects a reduction in power after removing samples from the longitudinally sampled females; FDR < 10%; 489 unique genes, as 25 genes were dominance-associated in multiple conditions, although the majority in the LPS-stimulated condition), 454 (88%) exhibited directionally consistent changes in mean expression levels across the dominance transition (Table S6). In other words, genes that were more highly expressed in dominant animals in the cross-sectional analysis also were more highly expressed in females that transitioned to dominant breeder status during the study, relative to the same females when they were subordinates. In comparison, in the control (matched sampling date) set, significantly fewer genes (60%) exhibited directionally consistent shifts (two sample proportion test, p = 5.5 x 10^-24^). Further, despite the small sample size, 95 of the 454 directionally consistent genes (21%) also exhibited large enough shifts to be identified as significant within-individual changes (p<0.05, paired t-test), and the dominance status effect size in the cross-sectional analysis is strongly correlated with the t-statistic from the longitudinal sample (Pearson r = 0.494, p = 6.88 x 10^-33^; Figure 4A). In the control set, only 17 (4%) of the same genes showed significant within-individual changes, consistent with the expected number of false positives across 454 tests (p<0.05, paired t-test). In addition, while there is a significant correlation between the dominance effect size from the cross-sectional analysis and the longitudinal sample t-statistic in the control set, it is much weaker than for females who transitioned to a dominant role (Pearson r = 0.133, p = 2.42 x 10^-3^; Figure 4B).

**Figure 4.**
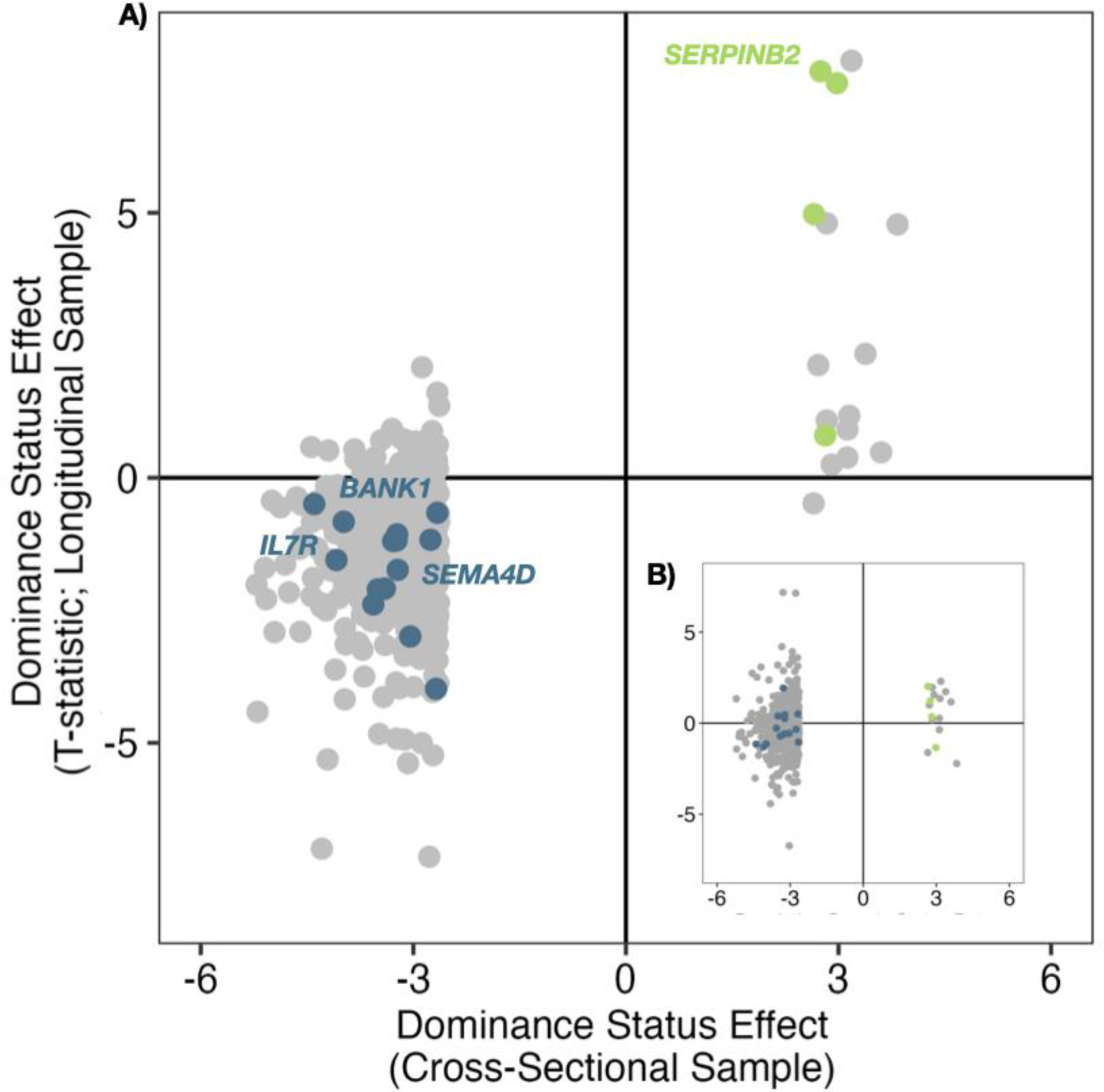
Status effects estimated from cross-sectional data are congruent with changes in gene expression in longitudinal transitions from subordinate to dominant. **A)** Standardized effect size of dominance status in females from cross-sectional analysis (x-axis: data from the LPS condition, where status effects are most apparent, are shown, based on analyses where only one sample was included from each set of longitudinal samples) are positively correlated with paired t-statistic values for the difference in gene expression between female meerkats when they transitioned from subordinate to dominant status (y-axis). Highlighted genes are members of key innate immune function gene sets enriched among status-associated genes. **B)** This positive correlation is not observed when calculating the parallel t-statistics for control meerkats sampled at the same time as those who transitioned from subordinate to dominant status (x and y axes are as in A).

### Cell composition and demographic correlates of female dominance do not explain the gene expression signature of breeding status

Dominant female meerkats are distinct from subordinate females in several key respects that do not define dominance *per se*, but could explain the observed gene expression signature of dominant status. Dominant females, including those in this study, tend to be heavier (dominant mean = 709.1 g ± 89.2 s.d.; subordinate mean = 626.5 g ± 81.6 s.d., Figure S1) and older (dominant mean = 3.13 yrs ± 1.54 s.d.; subordinate mean = 1.61 yrs ± 0.60 s.d., Figure S1; see also Clutton-Brock et al. (2006); Kutsukake and Clutton-Brock (2005)). They are also more likely to be sampled when pregnant: in our sample, for example, 9 of 15 captures of pregnant females were dominants, and only one of the pregnancies from subordinate captures resulted in live offspring (compared to 7 of 9 in dominants). In addition to controlling for pregnancy, body mass, and age in our main model, *post hoc* analyses showed that regressing out each of these variables from the gene expression data did not alter our findings for dominance status (r for dominance effect betas in the original model versus after first removing body mass, pregnancy, and age effects > 0.99, Figure S7).

Another possibility is that dominant and subordinate females differ in the composition of their peripheral blood mononuclear cells: that is, the signature of dominance is primarily a signature of compositional differences in the peripheral blood. The absence of meerkat-specific antibodies to markers of cell identity prevented us from profiling cell composition directly using flow cytometry. As an alternative, we estimated the composition of B cells, T cells, and monocytes in our sample based on the gene expression deconvolution method implemented in CIBERSORT (Newman et al., 2015, see Supplementary Methods). We found that dominant females tend to have slightly lower estimated fractions of T cells (t-test T statistic = 2.09, p = 0.04) and somewhat higher estimated fractions of B cells (T = -2.25, p = 0.03). However, including the T cell and B cell estimates in our differential expression models did not quantitatively alter our findings for social status (Figure S8, Pearson R, status effect sizes with and without T cell and B cell estimates, R=0.98-0.99, p < 0.001 across four challenge datasets). Further, while marker genes for T cells and B cells are enriched among genes that are associated with social status in female meerkats (FET adjusted p_Tcell_=5.08 x 10^-7^; p_Bcell_=1.7 x 10^-3^), genes in these marker sets (i.e., those for which high expression is a marker of a given cell type) are not more likely to be systematically upregulated or downregulated with dominant status (FET adjusted p_Tcell_=1; p_earlyBcell_=1; p_lateBcell_=1). Consequently, while we cannot rule out a contribution of status-related differences in cell composition to our results, they do not appear to make a major contribution to the signature of social status in our sample.

### Social status and gene regulation in social mammals

Social status predicts variation in gene regulation in birds, fish, insects, and mammals (e.g., Bondar et al., 2018; Lea et al., 2018; Lee et al., 2022a; Miller et al., 2009; Murray et al., 2019; Rittschof et al., 2014; Snyder-Mackler et al., 2016; Tung et al., 2012). To place our findings for meerkats in context, we therefore compared them to the gene expression signature of social status in two other social mammals that also profiled white blood cells in comparable control and LPS-challenged conditions: wild baboons from a majority yellow baboon (*Papio cynocephalus*) population in Kenya (male and female, Anderson et al. (2022)) and captive female rhesus macaques (*Macaca mulatta*: Snyder-Mackler et al. (2016)). Unlike meerkats, where there is only a single dominant animal of each sex in each group, and all other group members are classified as subordinate, both baboons and rhesus macaques are plural breeders that organize into linear, sex-specific hierarchies. Hierarchies are determined by nepotistic (matrilineal) inheritance of status in female baboons and rhesus macaques but by physical competition in male baboons.

In all three of these data sets, genes associated with social status are enriched in innate immune pathways (Figure 5A, Table S7). Incorporating the current findings into this comparison shows that the signature of social status in meerkat females more closely resembles the pattern for male baboons than the pattern for either female baboons or female rhesus macaques (Figure 5A). Like high-ranking male baboons, high status female meerkats tend to upregulate genes involved in TNFA signaling via NFkB and the interferon gamma response. In contrast, genes in the same pathways tend to be downregulated in high-ranking female macaques and female baboons (Table S7). Consequently, the gene-level effect sizes of dominance status are positively correlated between female meerkats and male baboons for genes involved in the inflammatory response, in an LPS-stimulated condition (Figure 5B, Pearson r = 0.261, p = 1.72 x 10^-2^). Gene-level effect sizes are also significantly negatively correlated between female meerkats and female baboons for genes involved in the complement system in the control condition (one of four pathways significantly enriched among dominance-associated genes in female meerkats: Figure 5C, Pearson r = -0.298, p = 5.53 x 10^-3^). Further, genes significantly associated with both dominance in female meerkats and dominance rank in male baboons tend to be directionally concordant (FET p = 3.26 x 10^-5^; the odds ratio cannot be calculated because no genes are dominance-associated in both cases and directionally discordant), while genes associated with both dominance in female meerkats and dominance rank in female macaques tend to be directionally *dis*cordant (FET log_2_(OR) = -2.75, p = 6.21 x 10^-4^). The direction of dominance effects in female meerkats did not predict the direction of dominance rank effects in female baboons (FET p > 0.05; the odds ratio cannot be calculated because few genes are associated with high status in both cases, and none of these genes are directionally concordant).

**Figure 5.**
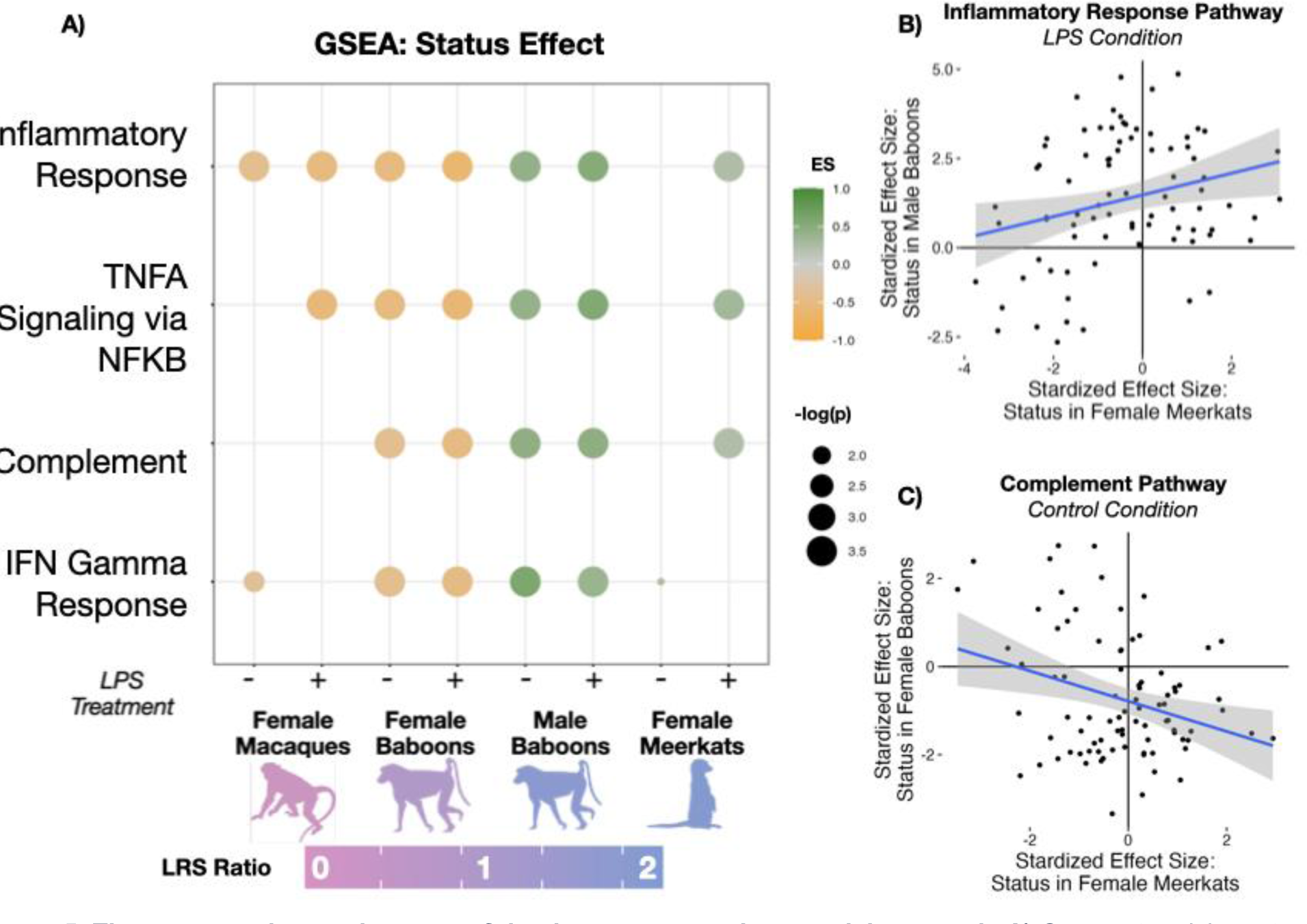
The gene regulatory signature of dominance across three social mammals. **A)** Gene set enrichment for immune pathways significantly enriched for dominance status effects in female meerkats (adjusted p-value < 0.05), across data sets. Green colors show pathways enriched among genes more highly expressed in high status individuals; orange colors show pathways enriched among genes more highly expressed in low status individuals. P-values are adjusted using Bonferroni correction based on running 27 total tests (all Hallmark gene sets in the immune, signaling, and pathways categories). One possible explanation for this difference relates to the intensity of investment in reproduction: colors for the silhouettes correspond to a bias towards greater reproductive skew in the focal sex than the opposite sex (purple), or to greater reproductive skew in the opposite sex than the focal sex (pink), based on estimates in Lukas and Clutton-Brock (2014). **B)** Gene-by-gene comparison of dominance status effects in female meerkats (x-axis; this study) and male baboons (y-axis; Anderson et al. (2022) for all measured genes in the inflammatory response pathway (r = 0.26, p = 0.017); **C)** Gene-by-gene comparison of dominance status effects in female meerkats (x-axis; this study) and female baboons (y-axis; Anderson et al. (2022) for all measured genes in the complement pathway (r = -0.30, p = 5.53 x 10^-3^).

Finally, we investigated concordance in status-related differences in Toll-like receptor 4 (TLR4) signaling. TLR4 is the primary receptor for LPS on monocytes and macrophages: LPS binding to TLR4 triggers signaling through alternative MyD88-dependent and TRIF-dependent pathways that stimulate NFkB-mediated and Type I interferon transcriptional cascades, respectively, and the relative use of these pathways has been linked to social status and/or social stress in both humans and other primates (Cole, 2014; Lea et al., 2018; Miller & Chen, 2007; Snyder-Mackler et al., 2016; Snyder-Mackler et al., 2019). In meerkats, MyD88-dependent genes that are upregulated during antigen stimulation (Ramsey et al., 2008) are strongly enriched among genes that are expressed more highly in dominant meerkat females and in response to LPS (FET log_2_(OR) = 3.42, p = 4.75 x 10^-3^). In this respect, too, female meerkats partially resemble male, but not female, baboons. Correlations between the effect of dominance rank in male baboons and dominant status in female meerkats among TRIF-dependent genes are significantly positive (r = 0.164, p = 0.012; MyD88-dependent genes showed a similar correlation, although not significant: r = 0.155, p = 0.055), but correlations between the effects of status in female baboons and female meerkats are significantly negative (MyD88 genes, r = -0.249, p = 2.04 x 10^-3^; TRIF genes, r = -0.159, p = 0.014). Thus, polarized TLR4 signaling after LPS challenge may be a consistent marker of social environmental conditions across species, but with variable directional effects.

## Discussion

Our results demonstrate a sex-specific signature of social status in immune cell gene expression in wild meerkats. This pattern extends previous findings in primates, in two species where males compete more intensely than females (Anderson et al., 2022; Lea et al., 2018; Snyder-Mackler et al., 2016), to a cooperatively breeding carnivore in which females compete more intensely than males (Clutton-Brock et al., 2001; Clutton-Brock et al., 2006; Clutton-Brock & Manser, 2016). Despite the substantial evolutionary distance between these taxa (carnivores diverged from primates more than 80 million years ago; dos Reis et al., 2012; Pozzi et al., 2014), several of the same pathways are associated with social status across lineages. These results suggest that a core set of genes is sensitive to differences in dominance status in the peripheral blood of social mammals.

However, the direction of these associations varies by species-sex combination. Like high-ranking male baboons, dominant female meerkats exhibit increased expression of innate immune defense genes and polarize the expression of genes in the TLR4 response pathway towards an NFkB-mediated pro-inflammatory response (Anderson et al., 2022). This pattern reverses that described for female baboons and female rhesus macaques. Consequently, our findings provide direct evidence that previously reported differences between male and female primates are not a consequence of sex *per se*, but instead arise due to correlates of social status that, depending on species, can characterize either males *or* females. One important correlate may be related to sex biases in reproductive skew, which is higher in male baboons than female baboons (and in male rhesus macaques versus female rhesus macaques, although we did not have gene expression data from male macaques here), but higher for female meerkats than male meerkats (Figure 5A; Alberts et al., 2006; Hodge et al., 2008; Lukas & Clutton-Brock, 2014; Spong et al., 2008; Widdig et al., 2004; Widdig et al., 2016). Notably, female-biased skew in meerkats is achieved through a combination of long tenure and high rates of pup production (Hodge et al., 2008), whereas male-biased skew in baboons is achieved through monopolizing mating opportunities with multiple females, typically over a shorter period of time (Alberts et al., 2003). An explanation based on reproductive skew would suggest that the gene expression signature of dominance in female meerkats and male baboons arises due to investment in reproductive competition, despite it manifesting differently between species.

In contrast to the other sex-species combinations evaluated here, we detected no measurable gene expression signature of dominance in meerkat males. Unlike in female meerkats, body mass does not predict the likelihood that male meerkats will attain dominant breeding status (instead, group size and composition are the primary predictor of reproductive success: (Spong et al., 2008). In this respect, they bear some resemblance to female macaques and baboons, where dominance rank is also largely independent of individual physical condition. However, unlike female primates or typical lab models for social defeat, dominance in male meerkats is not regularly reinforced by targeted harassment of subordinates. Subordinate males voluntarily seek out reproductive opportunities outside their natal groups and are seldom evicted (Stephens et al., 2005). One common explanation for status-related gene regulatory differences in social mammal hierarchies involves the chronic stress of social subordinacy, arising through systematic, directed agonistic behavior by higher-ranking individuals towards lower-ranking individuals (Anderson et al., 2022; Lea et al., 2018; Lee et al., 2022a; Lee et al., 2022c; Massart et al., 2017; Sanz et al., 2020; Snyder-Mackler et al., 2016; Tung et al., 2012). If neither testosterone levels, physical competition, nor regular received aggression differentiate dominant male meerkats from subordinate males, this may explain why they are also undifferentiable from the perspective of gene expression. For example, in cooperatively breeding wolves, where aggression from breeding animals to subordinates is similarly rare, gene expression signatures of dominance are also undetectable (Charruau et al., 2016).

Together, the emerging picture suggests that there are three dimensions of status hierarchies that may contribute to the presence and direction of associations between with social status and gene expression: first, the degree to which hierarchies are determined by physical competition and body condition; second, whether they result in high versus low reproductive skew; and third, whether status hierarchies are egalitarian or are regularly and systematically enforced. By focusing the present study on meerkats, we were able to disentangle these components of hierarchy dynamics from sex *per se*. However, quantifying the relative importance of physical competition during rank attainment and maintenance, energetic investment in reproductive opportunities, and behavioral enforcement of dominance will require further systematic exploration, including an expanded set of species-sex combinations. Such work may also benefit from investigating whether social status signatures are elevated during periods of intense competition for status (e.g., because of a recent dominance turnover or after immigration of new challengers; Alberts et al., 1992; Engh et al., 2006; Kaplan et al., 2009; Sapolsky, 1992) or attenuated in contracepted animals. Experimental manipulations could also help establish the degree to which status associations with gene regulation are *consequences* of rank-related differences in experience, versus a reflection of competitive ability (or other individual characteristics linked to rank attainment).

The present study leaves the proximate drivers and consequences of social status-associated gene expression differences unclear. Our analyses suggest that these consequences cannot be easily explained by differences in body mass, age, pregnancy status, or cell type composition (although compositional effects could occur at more granular levels of cell classifications than we were able to achieve here). A candidate behavioral mechanism is the rate of aggressive behavior, which mediates status-related variation in glucocorticoid levels in male chimpanzees (Muller et al., 2021) and social gradients in gene expression in captive rhesus macaques (Simons et al. (2022); but see Lea et al. (2018) in male baboons). Denser behavioral data or targeted sampling during subordinate evictions will be needed to test whether a similar phenomenon applies in female meerkats. At a physiological level, steroid hormones may also play an important role, as both glucocorticoid and testosterone levels are higher in breeding female meerkats than in subordinate females in this population (Barrette et al., 2012; Davies et al., 2016; Drea & Davies, 2022). Upon activation by hormone binding, glucocorticoid receptor and androgen receptor translocate into cell nuclei to remodel gene expression. While the evidence to date that steroid hormone levels explain status-related gene expression in social mammals is largely correlational, experimental manipulations of both androgen levels and glucocorticoid levels have been successfully conducted in wild meerkats, with consequences for cooperative and dominance-related behaviors (Dantzer et al., 2017b; Drea & Davies, 2022). The meerkat system thus has the potential to reveal how social status, endocrinology, and immune gene regulation are causally linked in a natural social mammal population, as a model for the social determinants of health and fitness more broadly.

Finally, the specific signature of dominance we observe in female meerkats points to elevated activity of inflammation-related pathways, a pattern that predicts morbidity and mortality in humans and that has been identified as a hallmark of aging (López-Otín et al., 2013). On the face of it, our results contradict the observation that dominant female meerkats in the Kalahari population live longer than subordinate helpers (Clutton-Brock et al., 2001). However, the greater longevity of dominant females, compared to subordinate females, is likely explained by increased extrinsic mortality risk to subordinates, who can be evicted and forced to spend more time separated from their social group in unfamiliar areas (Cram et al., 2018). Indeed, longitudinal sampling reveals that dominant female meerkats exhibit *faster* rates of telomere attrition, a biomarker of cellular senescence, than subordinates, despite their longer lifespans (Cram et al., 2017; Maag et al., 2022). Combined with our findings here, the emerging picture suggests that dominant meerkat females do experience physiological costs as a result of their repeated investment in reproduction, despite their longer mean lifespans (Thorley et al., 2020). Taken together, our results thus show that while studies of life history and demography alone have revealed examples of these trade-offs at an organismal level (e.g., Bauch et al., 2013; Fairlie et al., 2016; Lemaitre et al., 2014; Nussey et al., 2008; Sharp & Clutton-Brock, 2011), physiological and molecular analyses can contribute to a more complete picture.

## Acknowledgments

We thank the volunteers and managers of the Kalahari Meerkat Project for their contributions to maintaining the study population, field site, and data; Jack Thorley and Chris Duncan for assistance with data extraction and interpretation; Laurie Johnson and Doli Borah for assistance with sample organization; Reena Debray, Rachel Johnston, and Duncan Mahon for contributions to field lab set-up and sampling; Tim Vink for his contributions to the meerkat database; and Dave Gaynor for expert management of the field site. We also thank Anne Dumaine for coordination of sample preparation at the University of Chicago. *Ethics*. Ethical and sample collection clearance for this project was granted by the Northern Cape Province Department of Environment and Nature Conservation of South Africa (FAUNA 0930/2022 and FAUNA 1020/2016) and the Animal Ethics Committee of the University of Pretoria (EC047-16 and NAS003/2022). All research followed the standards outlined in the ASAB/ABS (2012) Guidelines for the Treatment of Animals in Behavioural Research and Teaching. *Funding.* We are grateful for funding support from the Human Frontier Science Program (RGP0051-2017 to JT, TCB, and LBB), the European Research Council Horizon 2020 initiative (ERC no. 742808 and 294494 to TCB), the Alfred P. Sloan Foundation (J.T.), the United States National Science Foundation (IOS-7801004 to J.T.), and the University of Zurich (M.M.). The long-term research on meerkats was supported by funding from the MAVA Foundation and the Universities of Zurich and Cambridge. Computational analyses were made possible by high-performance computing equipment funded in part by the North Carolina Biotechnology Center (2016-IDG-1013 and 2020-IIG-2109). CRC was supported in part through the United States National Institutes of Health F32AG067704.

## Data Accessibility

The raw reads generated in this study and a matrix of gene counts are available in the NCBI Gene Expression Omnibus (GEO) repository, GSE247525. Other data sets used in this study are contained within the article supplementary data files and analysis code is available at https://github.com/cryancampbell/meerkatPaper.

## Author Contributions

CRC, LBB, TCB, and JT conceived and designed the study. KW, MS, LBB, TCB, MM and JT collected data. CRC processed data and performed analyses. TCB, MM, LBB, and JT provided data and resources, provided supervision and oversight, and acquired funding. CRC and JT wrote the manuscript with edits and revisions from all authors.

## Notes

### Competing Interest Statement

The authors have declared no competing interest.

