## Supplemental Methods and Figures for "A female-biased gene expression signature of dominance in cooperatively breeding meerkats"

#### Supplemental Information

#### Table of Contents

**Supplementary Methods**
S1: Full linear model description
S2: Analysis of longitudinal samples
S3: Dominance-dependent polarization of TLR4 signaling
S4: Cell composition analysis with CIBERSORT

**Supplementary Figures**
S1: Correlates of dominance status in female meerkats
S2: TM 3'-seq libraries produce the expected mapping bias to the 3' ends of genes
S3: Association between pregnancy status and gene expression
S4: Gene expression responses to treatment by sex
S5: Gene expression associations with dominance by sex
S6: Enrichment of dominance associations with gene expression in females by treatment condition
S7: Stability of dominance effect estimates in females controlling for body mass, age, and pregnancy status
S8: Stability of dominance effect estimates in females controlling for CIBERSORT estimates of T cell and B cell proportions

**Tables** *(Campbell_etal_SupplementalTables.xlsx)*
S1: Capture metadata
S2: Sequencing metadata
S3 Model results: Treatment-control models
S4 Model results: All conditions, sex-specific
S5 Model results: Condition-specific, female-only
S6 Model results: Status x treatment interactions, female-only
S7 Longitudinal analysis results
S8 Multispecies GSEA results

### Methods S1: Full linear model description

To identify genes that were significantly associated with treatment condition or dominance status we used linear mixed effects models that control for relatedness within the sample, implemented in EMMREML (R package: EMMREML version 3.1, Akdemir & Okeke, 2015).

##### *i)* Treatment-control Models

We first explored the effect of a single treatment (LPS, Gard, or Dex stimulation) compared to control, both in the full data set (including both males and females) and in models for males and females separately. For each gene, for data from a single treatment (LPS, Gard, or Dex) with matched control, we estimated the effect of treatment condition and social status on gene expression levels using the following model:

*All Animals (Males & Females)*

$$(1)\quad y_{ijk} = \mu + t_j\delta + g_i\rho + s_{ik}\beta + a_{ik}\gamma + w_{ik}\eta + p_{ik}\nu + Zu + \varepsilon_{ijk},$$
$$u \sim MVN(0, \sigma_u^2 K), \qquad \varepsilon \sim MVN(0, \sigma_e^2 I)$$

where y is an n x 1 vector of gene expression levels across all sex-condition combinations, and $y_{ijk}$ is the residual gene expression level for individual *i* in treatment condition *j* for capture *k* (as some individuals were repeatedly sampled). $\mu$ is the intercept; *t* is an *n* x 1 vector of sample condition and δ is its effect size; *g* is an *n* x 1 vector of sex (female = 0, male = 1) and ρ its effect size; *s* is an *n* x 1 vector of dominance status (dominant = 1; subordinate = 0) and $\beta$ its effect size; *a* is an *n* x 1 vector of age in years at the time of sampling (centered) and $\gamma$ is its effect size; *w* is an *n* x 1 vector of individual body mass (centered), and $\eta$ is its effect size; and *p* is an *n* x 1 vector of pregnancy status (males = 0, females = -1 or 1), and $\nu$ is its effect size.

The vector *u* is an *m* x 1 vector is a random effects term to control for genetic structure in the population. Here, *m* is the number of unique animals in the analysis, the *m* x *m* matrix *K* contains estimates of pairwise relatedness derived from a pedigree (pedantics v 1.7, Morrissey & Wilson, 2010), $\sigma_u^2$ is the genetic variance component, and *Z* is an *n* x *m* matrix of 1's and 0's which maps gene expression measurements to unique individuals in *u* to account for repeated samplings from the same animals. Residual errors are represented by the n x 1 vector ε representing the environmental variance component, *I* is the identity matrix, and *MVN* denotes the multivariate normal distribution. The relatedness matrix is derived from pedigrees generated by genotyping 18 microsatellite loci (Leclaire et al., 2013).

To test for responses to the treatment condition, for each gene, we tested the null hypothesis that δ = 0 versus the alternative hypothesis, δ ≠ 0. To investigate dominance status effects, we tested the null hypothesis that $\beta = 0$ versus the alternative hypothesis, $\beta \neq 0$. We created empirical null distributions by permuting either treatment condition within capture or dominance status across captures, in blocks corresponding to treatment conditions for a single capture, 100 times and rerunning the analysis for each of the permutations. We then calculated false discovery rates using custom code (see: https://github.com/cryancampbell/meerkatPaper) that utilizes permutations following Storey and Tibshirani (2003).

We also performed parallel analyses to investigate responses to treatment within sex. These models were identical to Model 1 above, but removed the fixed effect of sex and were fit separately to either data for females only or data for males only.

##### *ii)* All conditions, sex-specific

We next investigated sex-specific associations with dominance rank by modeling data from all treatment conditions together, for either females only or males only, as follows:

$$(2)\ y_{ijk} = \mu + \delta_j t_j + s_{ik}\beta + a_{ik}\gamma + w_{ik}\eta + p_{ik}\nu + Zu + \varepsilon_{ijk},$$
$$u \sim MVN(0, \sigma_u^2 K),\ \ \varepsilon \sim MVN(0, \sigma_e^2 I)$$

where y is a n x 1 vector of gene expression levels across all conditions (control, dex, gard, and LPS), and $y_{ijk}$ is the residual gene expression level for individual *i* in treatment condition *j* for capture *k*. $\mu$ is the intercept; *t* is an *n* x 1 vector of sample condition (Dex = 1, Gard = 2, LPS = 3) and $\delta_j$ is the effect size (where control is the reference condition); *s* is an *n* x 1 vector of social status (dominant = 1; subordinate = 0) and $\beta$ its effect size; *a* is an *n* x 1 vector of age in years at the time of sampling (centered) and $\gamma$ is its effect size; *w* is an *n* x 1 vector of individual body mass (centered), and $\eta$ is its effect size; and *p* is an *n* x 1 vector of pregnancy status (-1 or 1), and $\nu$ is its effect size.

##### *iii)* Condition-specific, female-only

To investigate the conditions that drive the association between dominance status and gene expression in females, we ran a third set of models that investigated dominance effects within each condition separately.

$$(3)\ y_{ik} = \mu + s_{ik}\beta + a_{ik}\gamma + w_{ik}\eta + p_{ik}\nu + Zu + \varepsilon_{ik},$$
$$u \sim MVN(0, \sigma_u^2 K), \qquad \varepsilon \sim MVN(0, \sigma_e^2 I)$$

These models parallel equation 2 above, but remove the fixed effect of treatment because they fit only data from a single treatment condition (control, dex, Gard, or LPS).

##### *iv)* Status x treatment interactions, female-only

Finally, to investigate whether dominance status influences the gene expression *response* to stimulation, we fit a final set of models, for the set of 735 genes with significant associations with dominance status in Model (3).

$$(4)\ y_{ijk} = \mu + s_{ik}\beta_0 \times f(t=0) + s_{ik}\beta_1 \times f(t=1) + a_{ik}\gamma + w_{ik}\eta + p_{ik}\nu + Zu + \varepsilon_{ijk},$$
$$u \sim MVN(0, \sigma_u^2 K), \qquad \varepsilon \sim MVN(0, \sigma_e^2 I)$$

where y is a n x 1 vector of gene expression levels across the control and one stimulated condition (either Dex, Gard, or LPS, depending on the condition in which a significant association with status was identified in Model 3; no genes were identified in the control condition only), and $y_{ijk}$ is the residual gene expression level for individual *i* in treatment condition *j* for capture *k. f* is an indicator variable for the treatment condition (Control = 0, Treatment = 1), *s* is an *n* x 1 vector of social status (dominant = 1; subordinate = 0), $\beta_0$ is the effect size of status in the control condition, and $\beta_1$ is the effect size of status in the relevant treatment condition. To measure the interaction between treatment and status variables we compared the effect sizes of status estimated within Treatment ($\beta_1$) and Control ($\beta_0$) conditions. Here, positive interaction interaction effects indicate an increased response to the treatment condition in dominant animals. P-values for interaction effects were calculated on the standardized interaction effect sizes, treating the standardized effect size as a t-statistic.

### Methods S2: Analysis of longitudinal samples

To measure the effect of the transition from subordinate to dominant status on meerkat gene expression we focused on females intentionally sampled both before and after they transitioned status. To assess the difference in gene expression across this transition, we measured the log-fold change difference in gene expression between a sample collected when the individual was dominant and a sample collected, in the same treatment condition, when she was subordinate. If the animal had more than one sample taken when dominant or when subordinate, a single sample was chosen at random. We required females to be in the same pregnancy state across subordinate and dominant samples. For example, we excluded a repeated sample pair from the longitudinal analysis if an animal was captured only as a non-pregnant subordinate and then only as a pregnant dominant.

To test for significant within-individual shifts in gene expression when females transitioned to dominant status, we used paired t-tests for genes that exhibited congruent directional changes in longitudinal and cross-sectional analyses. To avoid biasing our comparison of cross-sectional and longitudinal analyses by including longitudinal data in the cross-sectional analysis, for all comparisons here we re-ran the cross-sectional analysis excluding all but one capture date for individuals used for the longitudinal comparison. Because we had fewer samples from dominant females than subordinate females, this exclusion meant, in practice, that we removed all samples collected when these females were subordinate. Importantly, this smaller data set recapitulates the results we obtain in the full data set, but with slightly reduced power (514 status-associated genes in instead of 709 in the full data set for Model 3, LPS condition).

### Methods S3: Dominance-dependent polarization of TLR4 signaling

To investigate status-dependent polarization of the TLR4 signaling pathways through TRIF and MyD88-dependent signaling pathways, we used the results from Ramsey et al (2008), who identified sets of genes in mouse for which the response to antigen induction was either MyD88- or TRIF-dependent. Ramsey et al compared the gene expression responses of macrophages from wild-type mice to those of MyD88 and TRIF knock-out mice, using six purified TLR agonists. They reported 334 genes that require MyD88 for normal stimulation-dependent increases in activity, and 274 that require TRIF. Of these genes, 126 and 103 have meerkat orthologues in our data set, respectively. Ramsey et al also reported 109 genes that require MyD88 for normal stimulation-dependent decreases in activity and 339 that require

TRIF, of which 48 and 155 have meerkat orthologues in our data set, respectively. Using these gene lists, we tested for enrichment of TRIF and MyD88-dependent genes among meerkat genes that are also upregulated in response to immune stimulation and are more highly expressed in dominant female meerkats. We also measured the correlation between standardized effect sizes for the effect of status in female meerkats with three species-sex combinations of non-human primate (female macaques, female baboons, male baboons) among all genes in the TRIF and MyD88-dependent gene sets.

#### Methods S4: Cell composition analysis with CIBERSORT

Using the Tabula Muris mouse single cell atlas (Schaum et al., 2018) we gathered marker information from droplet-sorted marrow cells. Schaum et al. (2018) determined marker genes for marrow cell subtypes including granulocytes, hematopoietic precursor cells, late pro-B cells, macrophages, early pro-B cells, monocytes, T cells, granulocytopoietic cells, erythroblasts, proerythroblasts, and basophils. In total, 53% of the marker genes for all subtypes had orthologues in the union set of all genes analyzed in meerkat models.

We used these gene markers as input for CIBERSORT (Newman et al., 2015), along with gene expression values in the control condition, to estimate the relative proportions of cells belonging to each subtype in the meerkat samples. In the initial run of CIBERSORT considering all 11 cell types from Schaum et al. (2018), several of the subtypes were assigned a proportion of 0 for the vast majority (>90%) of meerkat samples. We therefore re-ran CIBERSORT including only late pro-B cells, macrophages, early pro-B cells, monocytes, and T cells. We used these estimates to test for cell composition differences in by meerkat status across our sample, using t-tests.

Among these cell types, only B cells (late pro-B cells) and T cells were significantly, albeit weakly, correlated with female dominance status. We therefore fit *post hoc* models of gene expression including the B cell (late pro-B cells) and T cell estimates as additional fixed effects. These models otherwise follow the structure in Equation 3 above.

### Supplementary Figures

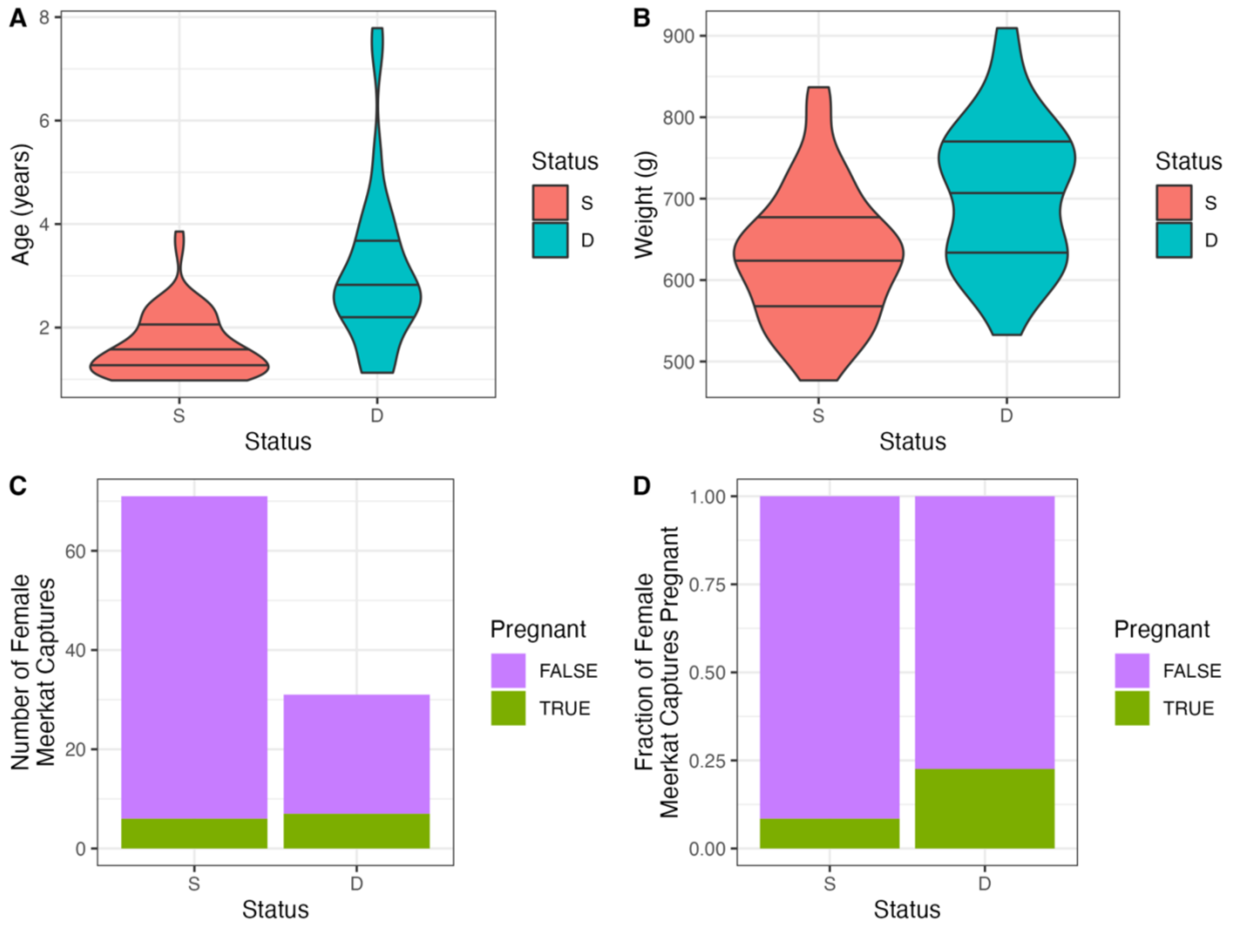


***Figure S1. Correlates of dominance status in female meerkats.*** *A) Violin plot comparing the ages at capture for dominant (D, blue) and subordinate (S, red) female meerkats (one-sided Wilcoxon Test, $W = 327.5$, $p = 9.5 \times 10^{-9}$). B) Violin plot comparing mean weight in the 60 days surrounding capture for dominant (D, blue) and subordinate (S, red) female meerkats (one-sided T-Test, $t = -4.18$, $p = 5.57 \times 10^{-5}$). C) Barplot displaying the number of dominant (D) and subordinate (S) animals that were pregnant (green) versus not pregnant (purple) at the time of capture (FET $\log_2(OR) = 2.58$, $p = 0.012$). D) Barplot displaying proportion of dominant (D) and subordinate (S) animals that were in early pregnancy (green) versus not pregnant (purple) at the time of capture.*

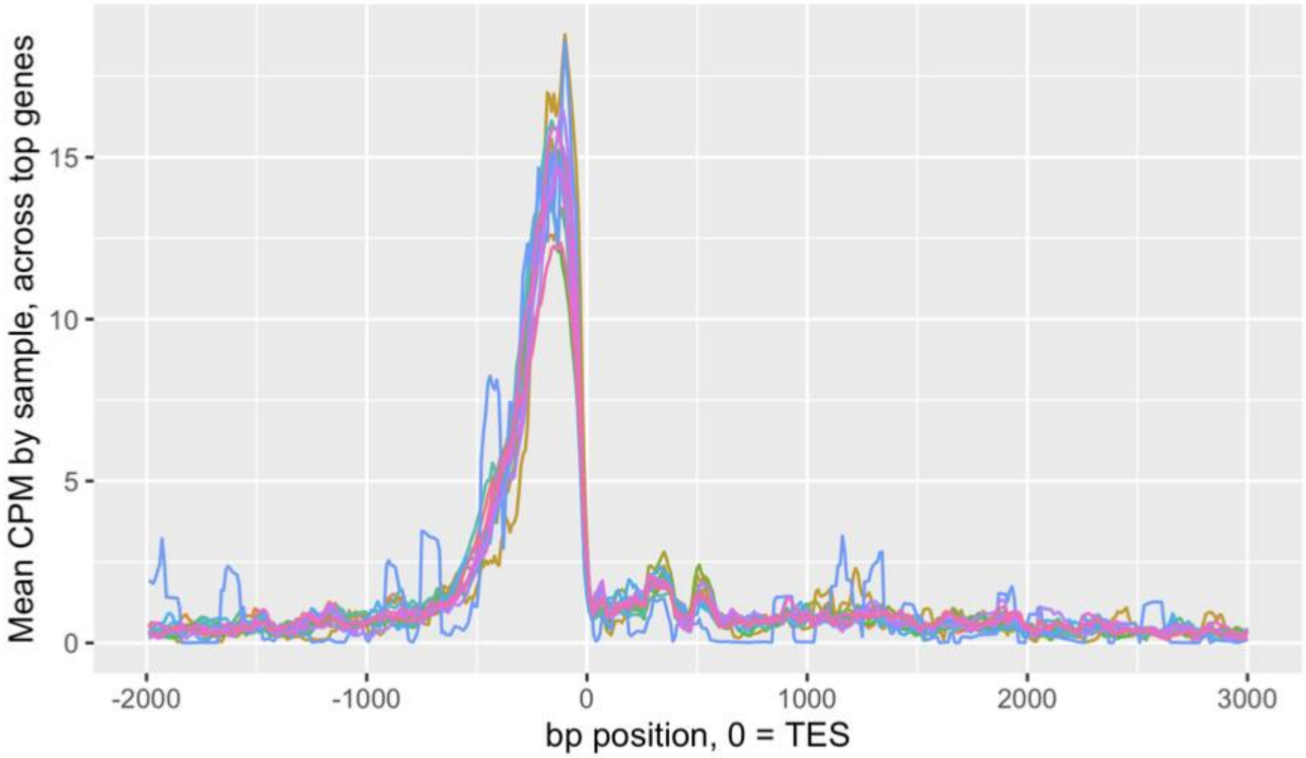


***Figure S2. TM 3'-seq libraries produce the expected mapping bias to the 3' ends of genes.*** *Mean CPM of top 8,141 expressed genes, plotted by position relative to transcription end site (TES), for a representative set of 15 meerkat samples.*

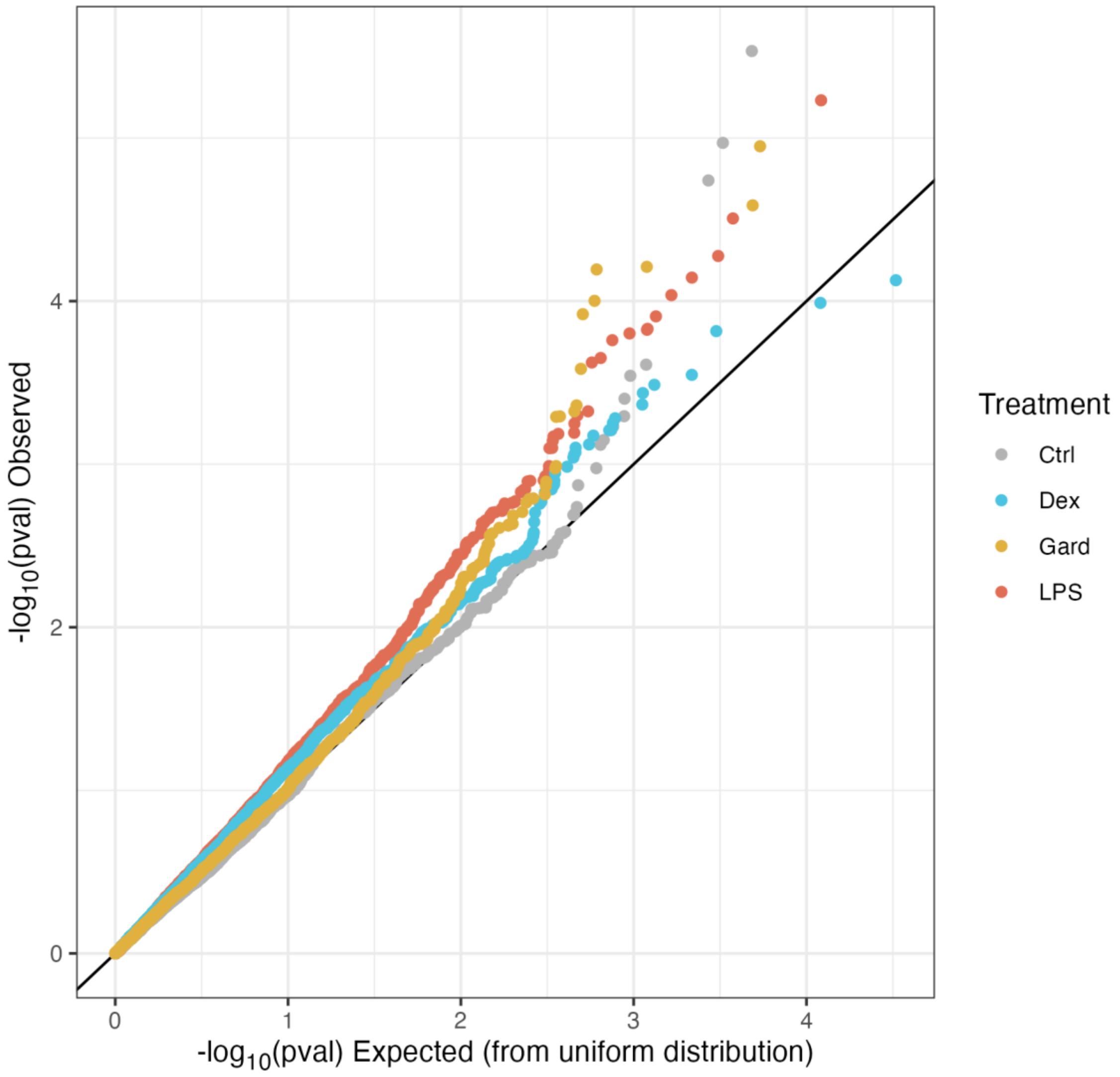


***Figure S3. Association between pregnancy status and gene expression.*** *Quantile-quantile plot of observed p-values (y-axis) versus expected p-values from a uniform distribution (x-axis), by condition (Control:gray, Dex:blue, Gard:yellow, LPS:red). The modest enrichment suggests a weak signature of pregnancy status in the gene expression data.*

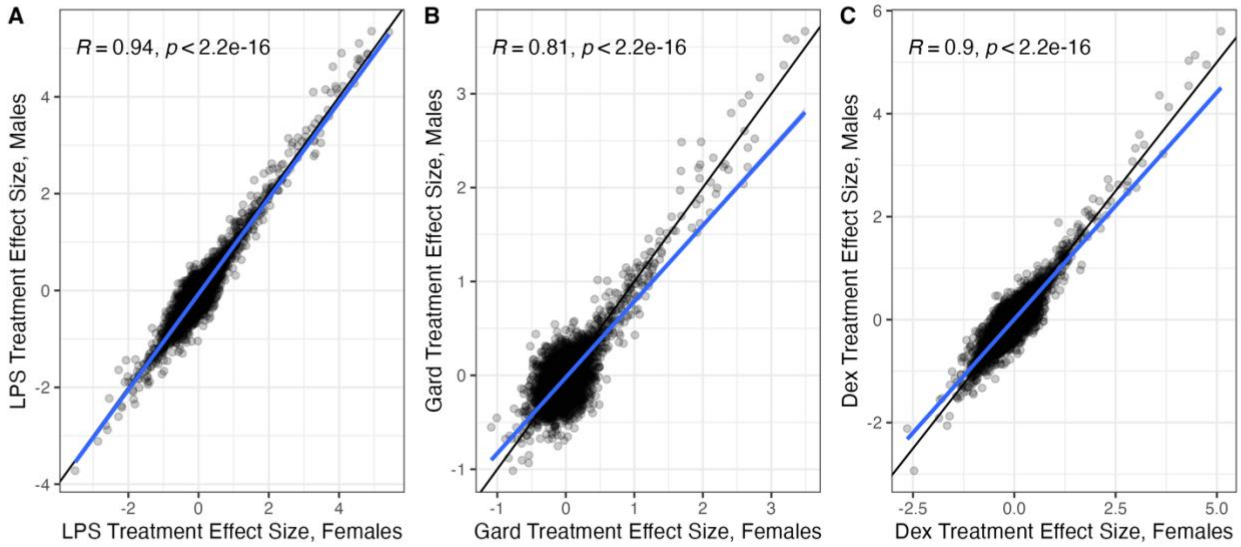


***Figure S4. Gene expression responses to treatment by sex.*** *The effect of treatment (A-LPS, B-Gardiquimod, C-Dexamethasone) on gene expression in male (y-axis) versus female (x-axis) meerkats. Black diagonal line shows x=y, and blue line represents line of best fit.*

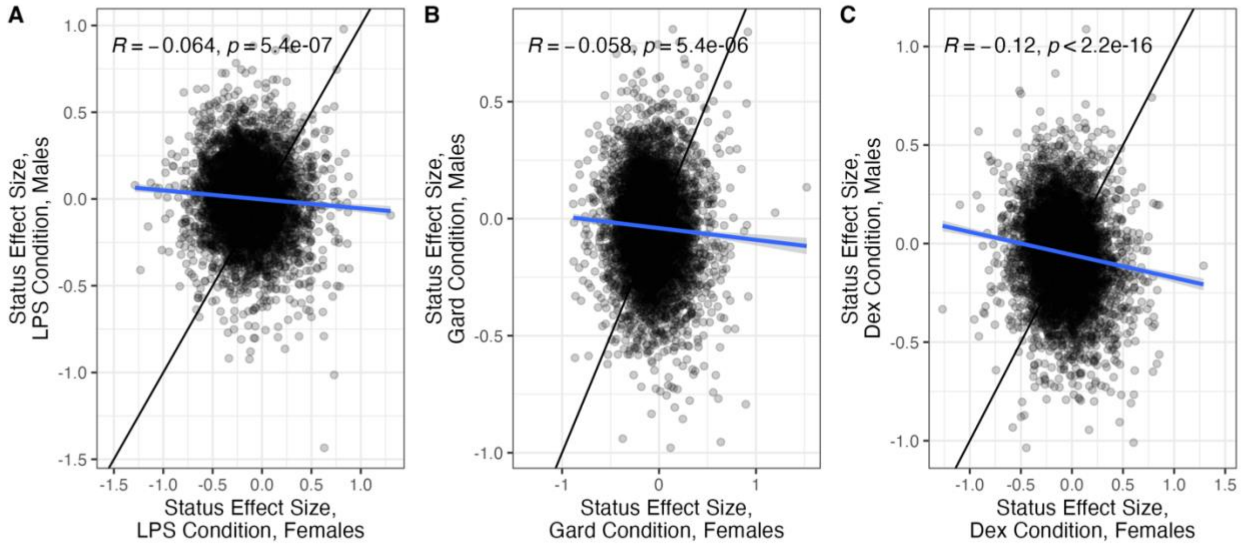


***Figure S5. Gene expression associations with dominance by sex.*** *The effects of dominance status in males and females are poorly correlated (A-LPS, B-Gardiquimod, C-Dexamethasone. Black diagonal line shows x=y, and blue line represents line of best fit.*

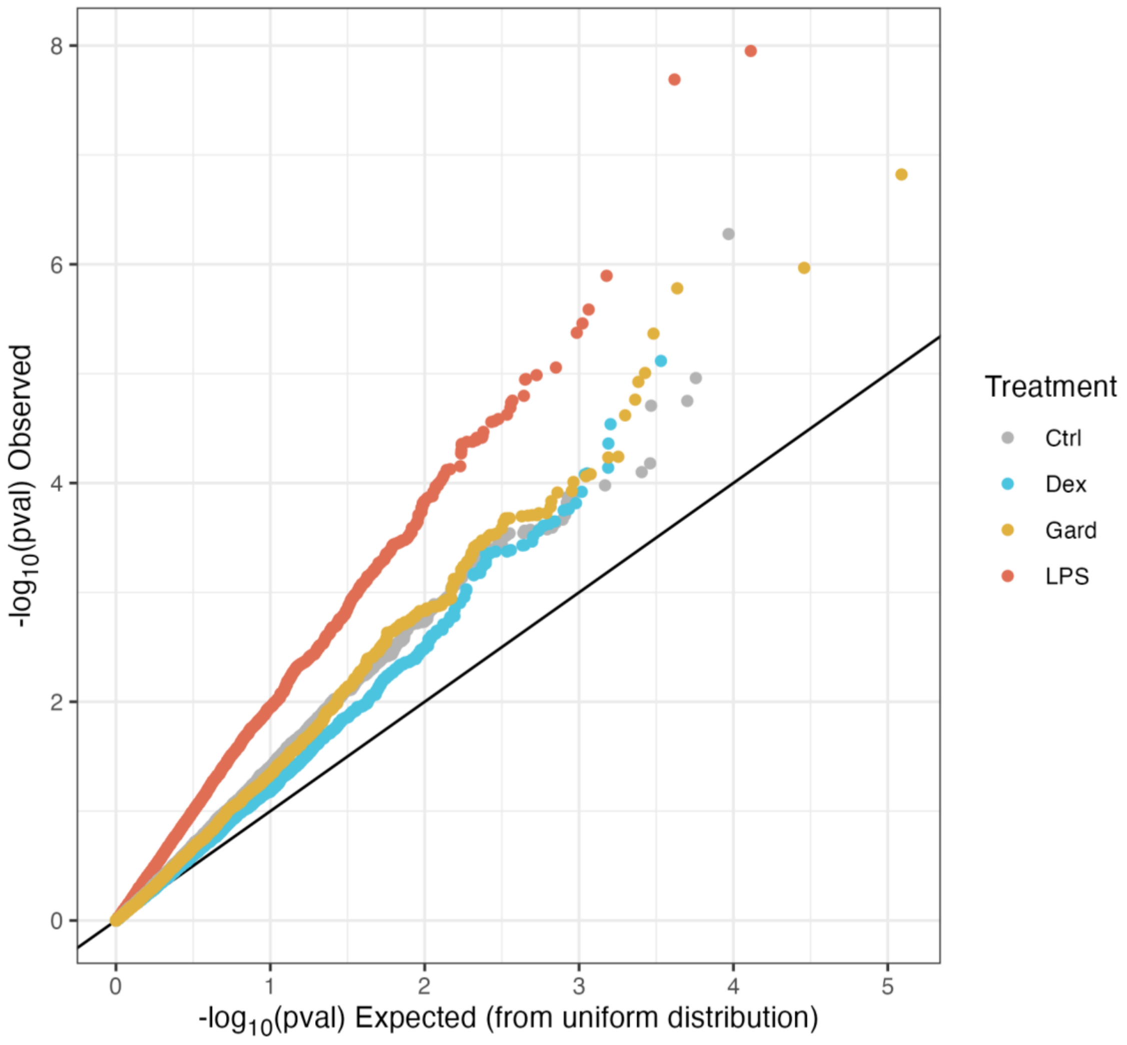


***Figure 6. Enrichment of dominance associations with gene expression in females by treatment condition.*** *Quantile-quantile plot of observed p-values (y-axis) versus expected p-values from a uniform distribution (x-axis), by condition (Control:gray, Dex:blue, Gard:yellow, LPS:red).*

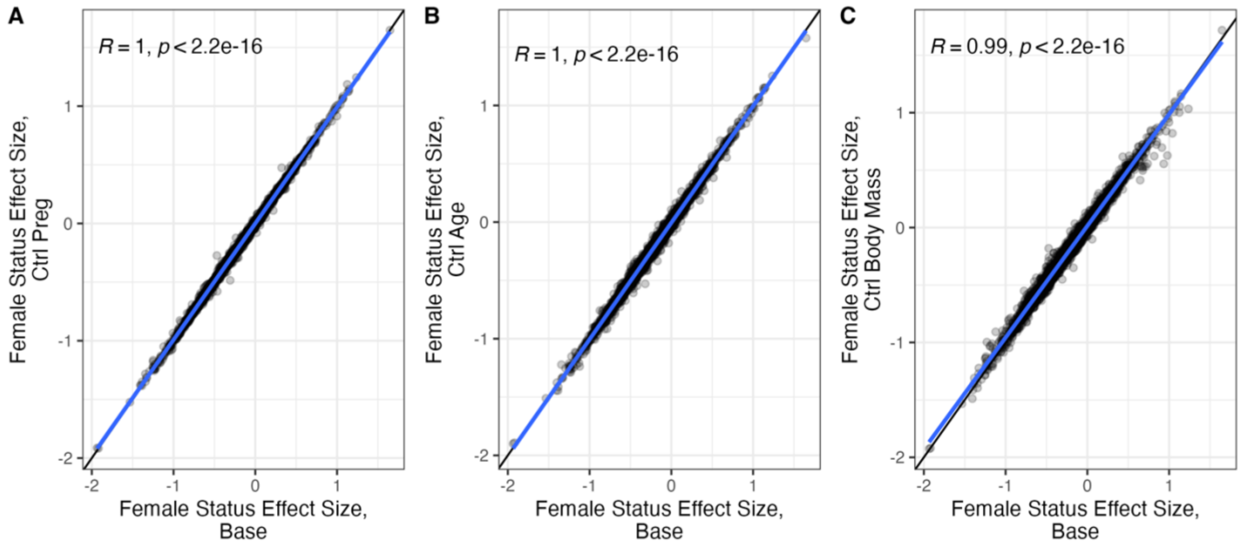


***Figure S7. Stability of dominance effect estimates in females controlling for body mass, age, and pregnancy status.*** *A) Estimated effect of dominance status in females, measured as in Model iii above (x-axis) versus estimated effect after regressing out pregnancy status from the gene expression data prior to modeling (y-axis). B) Estimated effect of dominance status in females, measured as in Model iii above (x-axis) versus estimated effect after regressing out age from the gene expression data prior to modeling (y-axis). C) Estimated effect of dominance status in females, measured as in Model iii above (x-axis) versus estimated effect after regressing out body mass from the gene expression data prior to modeling (y-axis).*

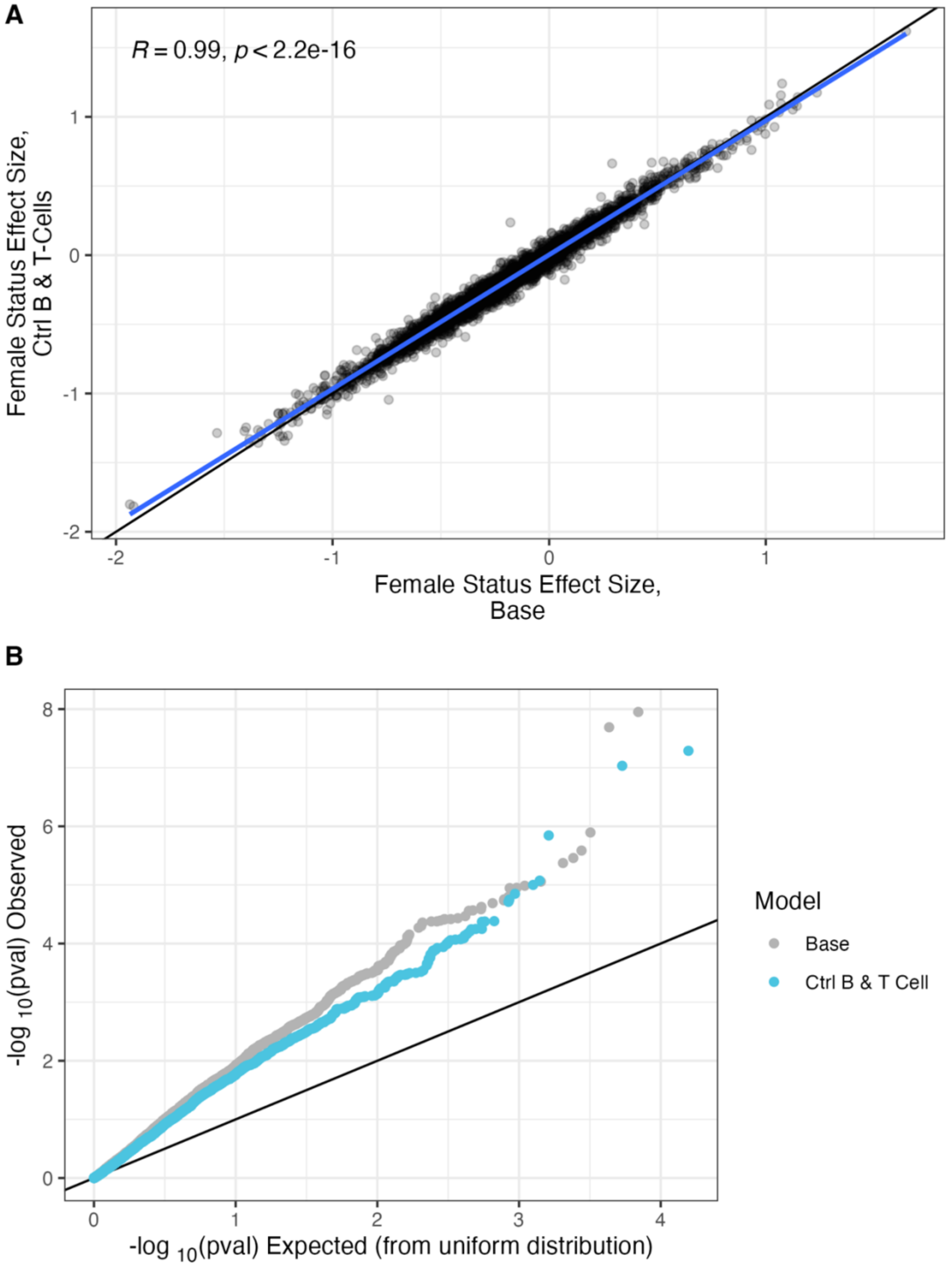


***Figure S8. Stability of dominance effect estimates in females controlling for CIBERSORT estimates of T cell and B cell proportions.*** *A) Estimated effect of dominance status in females as reported in Model iii (x-axis) versus when including estimates of B cell and T cell composition as fixed effect covariates (y-axis). Blue line represents line of best fit. B) qqPlot of the female dominance status effect, comparing observed p-values to those expected under a uniform null distribution. Gray dots show the qq plot for effects estimated in Model iii; blue dots show the same values when also controlling for cell composition.*
